# Bmp9 regulates Notch signaling and the temporal dynamics of angiogenesis via Lunatic Fringe

**DOI:** 10.1101/2023.09.25.557123

**Authors:** Tommaso Ristori, Raphael Thuret, Erika Hooker, Peter Quicke, Kevin Lanthier, Kalonji Ntumba, Irene M. Aspalter, Marina Uroz, Shane P. Herbert, Christopher S. Chen, Bruno Larrivée, Katie Bentley

## Abstract

**In brief:** The mechanisms regulating the signaling pathways involved in angiogenesis are not fully known. Ristori et al. show that Lunatic Fringe (LFng) mediates the crosstalk between Bone Morphogenic Protein 9 (Bmp9) and Notch signaling, thereby regulating the endothelial cell behavior and temporal dynamics of their identity during sprouting angiogenesis.

**Highlights:**

- Bmp9 upregulates the expression of LFng in endothelial cells.
- LFng regulates the temporal dynamics of tip/stalk selection and rearrangement.
- LFng indicated to play a role in hereditary hemorrhagic telangiectasia.
- Bmp9 and LFng mediate the endothelial cell-pericyte crosstalk.

Bone Morphogenic Protein 9 (Bmp9), whose signaling through Activin receptor-like kinase 1 (Alk1) is involved in several diseases, has been shown to independently activate Notch target genes in an additive fashion with canonical Notch signaling. Here, by integrating predictive computational modeling validated with experiments, we uncover that Bmp9 upregulates Lunatic Fringe (LFng) in endothelial cells (ECs), and thereby also regulates Notch activity in an inter-dependent, multiplicative fashion. Specifically, the Bmp9-upregulated LFng enhances Notch receptor activity creating a much stronger effect when Dll4 ligands are also present. During sprouting, this LFng regulation alters vessel branching by modulating the timing of EC phenotype selection and rearrangement. Our results further indicate that LFng can play a role in Bmp9-related diseases and in pericyte-driven vessel stabilization, since we find LFng contributes to Jag1 upregulation in Bmp9-stimulated ECs; thus, Bmp9-upregulated LFng results in not only enhanced EC Dll4-Notch1 activation, but also Jag1-Notch3 activation in pericytes.

## Introduction

The formation of new blood vessels from pre-existing ones, termed angiogenesis, is a complex biological process crucial in tissue development and healing. Dysregulated angiogenesis is correlated with several diseases such as cancer and hereditary hemorrhagic telangiectasia (HHT) (Carmeliet, 2003; Ardelean and Letarte, 2015). In particular, HHT is characterized by incorrect development of blood vessels and formation of arteriovenous malformations (AVMs), which can be caused by mutations of the activin receptor-like kinase 1 (Alk1). This receptor, together with its ligand Bone morphogenic protein 9 (Bmp9) (David *et al*., 2007; Chen *et al*., 2013), has been shown to be involved in the regulation of angiogenesis and vascular density (David *et al*., 2008; Larrivée *et al*., 2012). However, the exact contribution of the Bmp9/Alk1 pathway to the regulatory mechanisms of angiogenesis is not fully clear yet.

It is well established that the crosstalk between vascular endothelial growth factor (Vegf) and Notch signaling (Fig. 1A) is a fundamental regulatory mechanism of blood vessel formation (Hellström *et al*., 2007; Leslie *et al*., 2007; Lobov *et al*., 2007; Suchting *et al*., 2007). At the onset of angiogenesis, Vegf released by nearby hypoxic tissue is detected by Vegf-receptors present on endothelial cells (ECs) (Gerhardt *et al*., 2003). This stimulates the EC migratory response (Gerhardt *et al*., 2003) as well as upregulation of the ligand Delta-like 4 (Dll4) (Liu *et al*., 2003; Lobov *et al*., 2007). The consequential Notch1 activation in neighboring ECs leads transcription factors to modulate their Vegf receptor expression (Fig. 1A, blue pathway), such that these Notch1-activated cells obtain a decreased ability to sense Vegf and activate their migratory behavior (Williams *et al*., 2006; Lobov *et al*., 2007; Suchting *et al*., 2007; Harrington *et al*., 2008; Benedito *et al*., 2012). This cycle repeats over time, determining the formation of a pattern of highly migratory and Dll4-expressing cells (termed “tip cells”), alternated with less migratory and Notch1-activated cells (termed “stalk cells”) (Gerhardt *et al*., 2003). This ensures that only a few leading tip cells emerge from the existing vessels, with regular spacing, while proliferating stalk cells contribute to elongate the sprout (Gerhardt *et al*., 2003). The determination of these tip/stalk identities is nevertheless dynamic, since ECs overtake each other at the tip of sprouts (“cell shuffling”), continuously competing for the tip position via Notch signaling (Jakobsson *et al*., 2010; Arima *et al*., 2011; Bentley *et al*., 2014).

**Figure 1:**
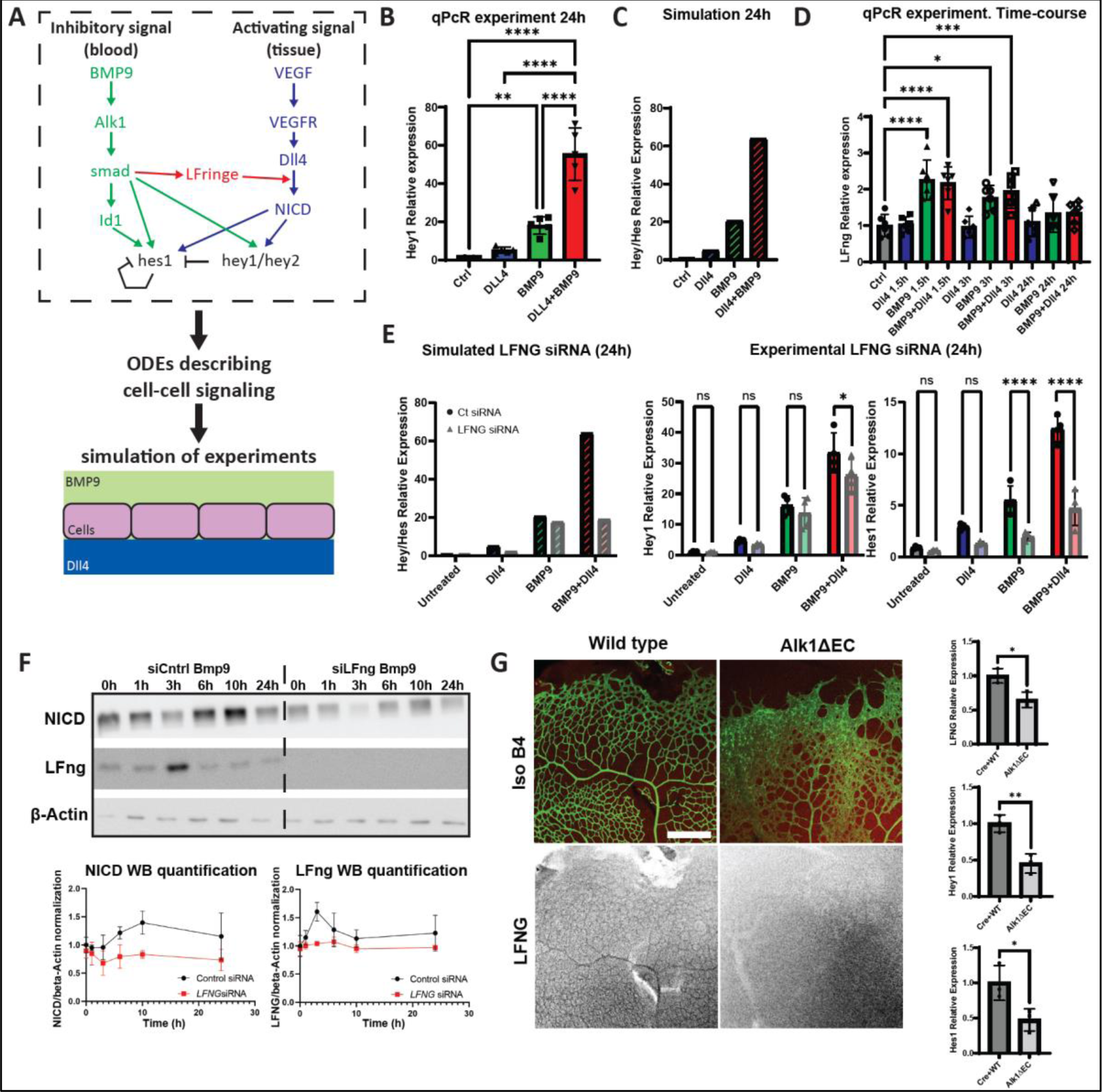
Bmp9 upregulates LFng expression in endothelial cells. (A) Schematic representation of the crosstalk between Bmp9 and Vegf-Notch signaling in ECs, as well as of the simulation and in vitro experiments. Green and blue lines represent previously established characteristics of the crosstalk; in red, the hypothesized LFng upregulation by Bmp9 is reported. (B-E) Experiments (full color) and simulations (striped color) of qPCR analysis of Hes1, Hey1 and LFng in HUVECs, induced by Bmp9 injection and/or Dll4 coating after 24h stimulation, with (bright color) or without (shaded color) LFng siRNA treatment. Number of repeats was higher than 4 in all cases ((B, D) N=5; (E) N=4). (F) Western blot analysis of NICD and LFng compared to the safe-keeping gene beta-actin in HUVECs with control (left) or LFng siRNA treatment (right) upon stimulation with 10ng/ml Bmp9, up to 24h, with relative quantification normalized over unstimulated data (bottom). The number of independent repeats per condition was N = 3. (G) IsolectinB4 and LFng staining of wild type and Alk1ΔEC retinal vessels, with associated normalized quantification of qPCR analysis. For the qPCR analysis, the number of independent experimental repeats per condition was N = 3. Scale bar, 200 µm. All values are mean +/− SD. *p < 0.05; **p<0.01; ***p<0.001; ****p<0.0001; (B, D-E) ANOVA, (G) Unpaired t-test.

Previous studies have shown that the activation of Alk1 by Bmp9 can independently activate Notch target genes to trigger stalk-cell markers and a consequential inhibition of angiogenesis (Larrivée *et al*., 2012; Moya *et al*., 2012). However, reported double-inhibition or gain-of-function studies on the Dll4-Notch and Bmp9-Alk1 pathways combined (Larrivée *et al*., 2012) demonstrated a multiplicative increase or decrease in angiogenic branching, respectively, that the prevailing additive, independent-activation of Notch target genes model (Fig. 1A, green pathway) of crosstalk between them cannot explain. Thus, we hypothesized that additional, inter-dependent crosstalk must exist between the two pathways, such that modulation of one can create positive feedback to enhance the effects of the other.

Here, we examined the possibility that the Bmp9 and Notch pathways might be linked by Fringe proteins, previously shown to be important for sprouting angiogenesis. Lunatic, Manic, and Radical Fringe are enzymes that increase Notch1 and Dll4 binding affinity by glycosylating the Notch extracellular domain. Lunatic Fringe (LFng) has a dominant role in the regulation of Notch1 signaling (Pennarubia *et al*., 2021), and its knockout is known to cause vascular hypersprouting resulting from Notch inhibition in ECs (Benedito *et al*., 2009). Despite its key role in angiogenesis, however, the regulators of LFng during angiogenesis are currently largely unclear. Previous studies have shown that LFng expression can be controlled by Notch activity in somites (Morales, Yasuda and Ish-Horowicz, 2002). In ECs, it has been shown that LFng is modulated by the endothelial transcription factor ERG (Shah *et al*., 2017). Interestingly, ERG also regulates the effects of Bmp9 on ECs (Dufton *et al*., 2017). Overall, these studies suggest a possible link between Bmp9 and LFng in ECs.

By integrating predictive computational simulations with *in vitro* and *in vivo* experiments, we identified that BMP9 upregulates LFng, which in turn enhances Notch signaling in an interdependent crosstalk manner and affects the temporal dynamics of Notch patterning. We previously showed that modulation of Notch pattern timing can delay or accelerate tip/stalk cell selection and shuffling, resulting in sparser or denser branch spacing during angiogenesis and thereby impacting the entire, resulting vascular network topology (Kur *et al*., 2016; Bentley and Chakravartula, 2017; Page *et al*., 2019; Zakirov *et al*., 2021). For example, a slower process of tip/stalk cell selection led to a sparser vascular network topology, e.g. when the anti-angiogenic, tissue derived factor semaphorin3E, which co-regulates Dll4, was lost (Kur *et al*., 2016). We have previously found via theoretical modeling that LFng alone could potentially modulate Notch pattern timing (Venkatraman, Regan and Bentley, 2016). Here, via extended modelling integrated with experiments, we demonstrate that Bmp9 regulation of LFng can alter Notch pattern timing, branching, and shuffling *in silico*, *in vitro* and *in vivo*. Simulations with an agent-based model, compared with additional *in vivo* experiments, also highlighted that the BMP9-mediated LFng upregulation can play a role in BMP9-related diseases, impacting cell clustering during AVM formation. Finally, we show that Bmp9-mediated upregulation of LFng potentiates the Notch crosstalk between pericytes and ECs, which can affect the vital pericyte-mediated stabilization of ECs at the end of the angiogenic process. The discovery of LFng as a strong enhancer of the Bmp9-Notch crosstalk and temporal modulation of tip selection and branch spacing, points to a new target for potential development of novel medical therapies aimed at regulating and improving vascularization in disease.

## Results

### Simulations predict Bmp9 upregulates a Fringe protein to enhance Notch ligand-receptor binding rate

We hypothesized that a missing, interdependent crosstalk link must exist between the Bmp9/Alk1 and Vegf/Notch pathways. This was motivated by previous qPCR analysis of human umbilical vein endothelial cells (HUVECs) exposed to either Bmp9, Dll4-coated substrates, or both, which highlighted a multiplicative increase upon double stimulation that cannot be explained by the prevailing (additive) independent crosstalk model (Fig. 1A, green-blue schematic) (Larrivée *et al*., 2012). This *in vitro* setting mimics the stimuli that ECs sense during angiogenesis, when they are exposed to paracrine Bmp9 and Dll4 present in neighboring ECs. Here, we first repeated those experiments and observed the same multiplicative effect (Fig. 1B). HUVECs upregulated the Hey1 expression in response to Dll4 (∼5-fold increase) and Bmp9 (∼18-fold increase) and, most interestingly, the upregulation resulting from the two stimuli combined (∼55-fold increase) was far greater than a simple addition of the effects resulting from the two stimuli taken alone (Fig. 1B).

To test our hypothesis and investigate the possible inter-dependent effects that Bmp9 could have on Notch signaling, we utilized computational modeling by extending our existing ordinary differential equation (ODE) model of Vegf/Notch signaling during angiogenesis (Venkatraman, Regan and Bentley, 2016). Previous studies validated this model’s predictive capability *in vivo* (Page *et al*., 2019). Briefly, the computational model assumes that external Vegf detected by Vegfr2 upregulates the formation of filopodia, which move receptors further into a Vegf gradient, creating positive feedback to Vegf activation and the expression of Dll4 (Zakirov *et al*., 2021), which can in turn bind to Notch1 expressed in neighboring cells. The consequential Notch1 activation upregulates the expression of target genes of Notch (e.g. Hey1 and Hes1, collectively termed HE in the model). This upregulation, in turn, downregulates Vegfr2 in the receiving, Notch activated neighboring cell. Here, we adapted the model to simulate cell signaling among a line of 10 to 50 cells (Fig. 1A), instead of only 2 cells as in the original study (Venkatraman, Regan and Bentley, 2016). This was performed by accounting for the movement of ligands across the cell membrane (see Methods section for details). The line of connected ECs was simulated as being exposed to Dll4 externally, mimicking the Dll4 coating as per the *in vitro* experimental conditions (Fig. 1A, (Larrivée *et al*., 2012)). The original model parameters were used for cell-cell signaling, while the level of Dll4 coating was calibrated to match the 5-fold increase in Hey1 expression as observed in the experiments (Fig. S1A, horizontal dotted lines). To explore how different upstream Bmp9-Notch crosstalk interactions would impact the Dll4-coating effects on Notch activation levels, each model parameter was either halved or doubled, to respectively mimic Bmp9-mediated downregulation or upregulation of the associated biological phenomenon. This parameter exploration predicted that the synergistic effects of Dll4 and Bmp9 on Hey1 can be best explained by an increase in the Dll4-Notch1 binding rate or the Notch1 expression, or by a decrease in the Notch1 or NICD degradation rates (Fig. S1A). Indeed, for example, by assuming that Bmp9 increases not only Hey1 expression, but also the Dll4-Notch1 binding rate (Fig. 1A), the simulations can now accurately replicate the otherwise unexplained experimental multiplicative result shown in Fig. 1B-C, when Dll4 and Bmp9 are co-modulated.

The binding rate of Notch ligands and receptors is positively correlated with their affinity (Luca *et al*., 2017). It is well known that the affinity of Notch1 with its ligands is strongly influenced by the enzymes Lunatic, Manic, and Radical Fringe (Kakuda and Haltiwanger, 2017; Kakuda *et al*., 2020). These enzymes have been shown to have a key role in the regulation of angiogenesis (Benedito *et al*., 2009), and previous modeling from our group showed that modulation of LFng might influence the temporal dynamics of tip-stalk selection, slowing tip selection down if lost (Venkatraman, Regan and Bentley, 2016). Therefore, we decided to focus on investigating whether the multiplicative effects of Bmp9 and Notch modulation on angiogenesis are mediated by Fringes experimentally.

### Bmp9 upregulates the expression of LFng *in vitro* and *in vivo*

As suggested by the computational findings, we next checked whether Bmp9 modulates the expression of the Fringe enzymes experimentally. HUVEC monolayers were first starved for 24h and then exposed to either Bmp9 added to the medium, Dll4-coating, or both; the resulting Fringe expression was then measured via qPCR. Considering that feedback loops could possibly cause temporal oscillations of the enzyme expression (Wöltje, Jabs and Fischer, 2015; Kur *et al*., 2016; Page *et al*., 2019), different time points were analyzed: 1.5h, 3h, and 24h. The qPCR analysis showed that, differently than Manic and Radical Fringe (Fig. S1B), Lunatic Fringe (LFng) is strongly and rapidly upregulated by Bmp9 after 1.5h, increasing at 3h, followed by a return to untreated levels after 24 hours (Fig. 1D). This upregulation was verified by Western Blotting (Fig. 1F), which confirmed at the protein level that the effects of Bmp9 on LFng are time-dependent and relatively rapid: LFng was upregulated and increased at 3h after Bmp9 exposure, while the effects decayed later. Therefore, our coupling between computational simulations and experiments uncovered that Bmp9 alters LFng expression in a time-dependent manner.

Western Blotting further confirmed that Bmp9 exposure leads to LFng- and Dll4-mediated Notch activation. In fact, Western Blotting for the Notch Intracellular Domain (NICD) indicates a strong increase in Notch cleavage at 6h and 10h, which is lost when LFng or Dll4 are reduced by small interfering RNA (siRNA) treatment (Figs. 1F, S1E). Finally, to validate Alk1 signaling regulation of LFng *in vivo*, the retina of wild type and Alk1ΔEC mice was stained for LFng. Consistent with our *in silico* and *in vitro* results, conditional heterozygous deletion of Alk1 in the endothelium caused a sharp decrease in the intensity of LFng staining (Fig. 1G), thereby confirming that Bmp9-mediated activation of Alk1 regulates LFng expression in ECs *in vivo*. This was confirmed by qPCR analysis of retinas, showing that LFng levels were significantly lower in Alk1ΔEC mice and, accordingly, Hes1 and Hey1 expressions were also decreased.

### LFng enhances Bmp9-mediated upregulation of endothelial Notch target genes

We next investigated whether LFng regulation by Bmp9 enhances activation of Notch target genes, alone and when in combination with Dll4 coating, to identify whether it can at least in part explain the multiplicative increase in target gene expression when both Dll4 and Bmp9 are co-modulated (Fig. 1B).

First, we adapted the ODE model to simulate a line of connected HUVECs exposed to Bmp9 (Figs. 1A, 1E), with and without LFng knockdown (KD). Given the influence of LFng on the receptor-ligand affinity (Kakuda and Haltiwanger, 2017) and thus on their binding rate (Luca *et al*., 2017), LFng KD was simulated by decreasing the Dll4-Notch1 binding rate. The simulations predicted that LFng KD decreases the effects of Bmp9 on the expression of Notch target genes Hes/Hey (HE in the model). The decrease was particularly evident in the presence of combined Bmp9 and Dll4 coating, for which the multiplicative effect was lost and the increase of Hes/Hey was almost exclusively due to the direct effect of Bmp9 on these target genes (Fig. 1E). These computational results therefore suggested that LFng mediates a proportion of the Bmp9 and Notch effects on ECs, especially enhancing them when Dll4 is present.

To verify these computational findings, we measured the expression of key Notch target genes in HUVECs *in vitro* exposed to Bmp9 with or without Dll4 coating, under control or LFng KD conditions. LFng KD HUVECs were obtained by transfecting these cells using LFng small interfering RNA (siRNA). This transfection downregulated LFng expression compared to control experiments, performed with control siRNA, with and without Bmp9 (Fig. S1C). In agreement with the simulations, the Bmp9-mediated upregulation of the Notch target genes Hes1/Hey1, observed in previous studies (Larrivée *et al*., 2012; Moya *et al*., 2012; Kerr *et al*., 2015), was significantly reduced by LFng siRNA exposure (Fig. 1E) when ECs were exposed to both Bmp9 and Dll4 coating, with approximately a 22% and a 63% reduction for Hey1 and Hes1, respectively. LFng KD significantly decreased Hes1 expression also when only Bmp9 was present, with approximately a 56% reduction, while this was not the case for Hey1. This is consistent with previous studies showing that, in synergy with cell serum, Bmp9 can induce Hey1 expression, but not Hes1, in a Notch-independent fashion (Wöltje, Jabs and Fischer, 2015). Overall, our results indicate that the LFng upregulation by Bmp9 mediates a significant proportion of the effects Bmp9 has on ECs.

### Bmp9 and LFng are temporal regulators of tip/stalk cell selection *in silico*

At the onset of angiogenesis, ECs exposed to Vegf communicate with their direct neighbor via juxtacrine Notch signaling, which determines the formation of a salt-and-pepper pattern of selected migratory tip cells alternated by Notch activated, less motile stalk cells (Hellström *et al*., 2007; Leslie *et al*., 2007; Lobov *et al*., 2007; Suchting *et al*., 2007). Previous experimental evidence demonstrated that the temporal dynamics of this tip/stalk cell selection strongly affects the resulting vascular topology (Kur *et al*., 2016; Ubezio *et al*., 2016; Bentley and Chakravartula, 2017), where slower tip selection leads to sparser branch spacing. Given the importance of these temporal dynamics on the resulting network topology, we next performed computational simulations to investigate whether Bmp9 and LFng are temporal regulators of tip selection. In the model, after simulated starvation (1 day without Vegf addition) an array of connected cells was exposed to Vegf, and the time to establish a tip/stalk “salt and pepper” pattern was quantified as in previous studies (Bentley, Gerhardt and Bates, 2008; Zakirov *et al*., 2021, see methods for more details) (Fig. 2A-B). We found that the time it took to establish a salt and pepper pattern was different depending on the conditions simulated (Figs. 2A-B, S2). For example, when cells were exposed to Bmp9 (in addition to Vegf), they retained the features of stalk cells for a longer duration, which slowed down the selection compared to control (Figs. 2A-B).

**Figure 2:**
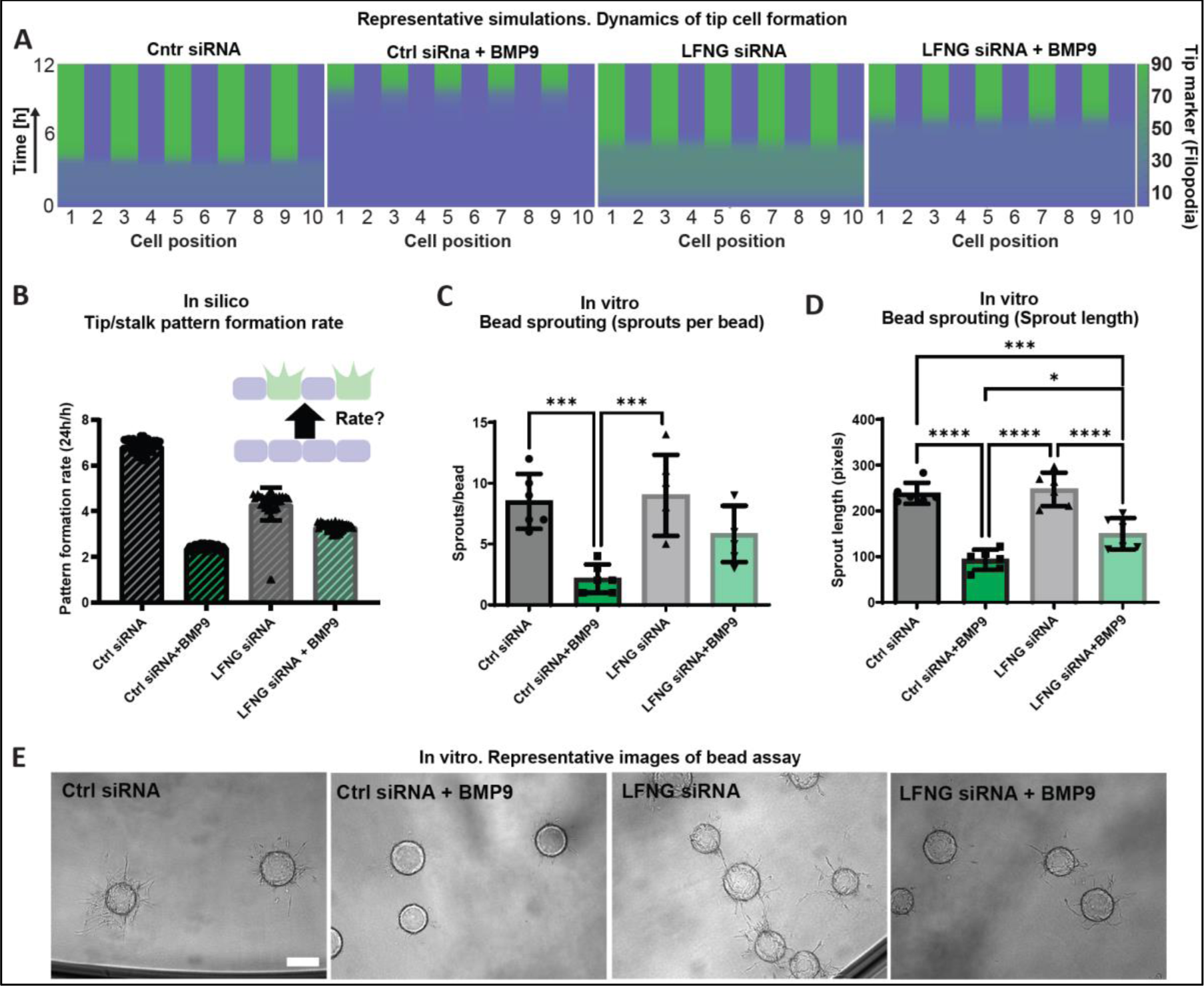
LFng mediates Bmp9 effects on angiogenesis. (A-B) Representative simulations of tip/stalk pattern formation for a row of 10 ECs exposed to Vegf with and without Bmp9 and LFng siRNA treatment and associated pattern formation rate. The color scheme in (A) represents the amount of filopodia in each cell. The pattern formation rate in (B) was approximated as the ratio between 24h at the nominator and the time necessary to pattern at the denominator. While Bmp9 slows down pattern formation, LFng siRNA treatment can partially rescue the control pattern formation rate. The number of simulations for each condition was N = 50. (C-E) Quantification (C-D) and representative images (E) of bead assays performed with HUVECs in fibrin gels, with or without Bmp9 stimulation (1 ng/ml) and LFng siRNA treatment. Bmp9 stimulation reduces the number of sprouts per bead (C) and the sprout length (D). LFng siRNA could significantly increase the sprout length in the case of Bmp9 stimulation, while a non-significant increase was observed for the number of beads per sprout. The number of independent experimental repeats per condition was N = 5. Scale bar, 100 µm. All values are mean +/− SD. *p < 0.05; **p<0.01; ***p<0.001; ****p<0.0001; ANOVA.

The simulations predict that LFng levels have biphasic effects on the time to pattern (Fig. S2A, S2E). Decreasing LFng caused a delay or even absence of the pattern formation, at least during the simulated time window of Vegf exposure (chosen as 24h to match the monolayer experiments). This occurs because Dll4-Notch1 binding is reduced when LFng is inhibited. Consequently, Notch1 activity is decreased, and all cells retain higher Vegfr2 and filopodia formation despite overall high levels of Dll4. This resulted in a large number of ECs rapidly becoming tip cells (Fig. 2A, S2A), but having a scarce efficiency to inhibit their neighbours, with a consequential absence or delay of the tip/stalk pattern formation. On the other hand, increasing LFng compared to the original value slightly slowed down pattern formation (Fig. S2E) because of the resulting generally high Notch activity, which decreased Vegfr2 levels and slowed down the formation of filopodia. In contrast, Hey1/Hes1 levels had a monotonous effect; increasing their level slowed down pattern formation (Fig. S2F). This occurred because Hes1/Hey1 directly downregulate Vegfr2 (Henderson *et al*., 2001) and, therefore, they slow down the detection of Vegf and the consequential filopodia formation and tip cell selection.

In our model, Bmp9 upregulates Hey1/Hes1 expression, both directly and via a LFng-mediated increase of Dll4-Notch binding activity. Consistent with the Bmp9-mediated LFng upregulation, the Bmp9 levels also exhibited a biphasic effect on the time to pattern, with an initial acceleration to the time to pattern and then a considerable deceleration after a minimum level was reached (Fig. S2G). In our simulations, LFng KD upon Bmp9 exposure could partially rescue the patterning speed, bringing it closer to the one observed with original parameters (Fig. 2B, S2H). These results can be explained by the fact that, with LFng inhibition, Bmp9 can enhance Hey1/Hes1 expression only directly, rather than also indirectly via Dll4-Notch binding changes. In conclusion, the computational simulations indicate that Bmp9 and LFng are temporal regulators of tip/stalk selection, slowing it down when Bmp9 is added, consistent with the sparser branching network topology observed *in vitro* and *in vivo* when Bmp9 is added (Larrivée *et al*., 2012; Moya *et al*., 2012).

### LFng mediates Bmp9 effects on *in vitro* angiogenic sprouting

To get a first verification that LFng mediates Bmp9 effects on sprouting angiogenesis, we performed a bead sprouting assay by using HUVECs that were previously transfected with LFng siRNA or control siRNA, with and without addition of Bmp9 (Fig. 2E). The samples were fixed and imaged to quantify the number of sprouts per bead and their length, as a quantification of the density of the vascular network. In agreement with previous *in vivo* studies (Larrivée *et al*., 2012) and the deceleration of tip/stalk pattern formation predicted by our simulations (Fig. 2B), addition of Bmp9 stimulation significantly decreased the length and number of sprouts compared to control (Fig. 2C-E).

Our computational results predicted that inhibiting LFng upon Bmp9 injection should partially rescue branching tip selection speeds, and therefore vascular density (Fig. 2B). Consistently, LFng downregulation was able to partially rescue branching; upon Bmp9 stimulation, LFng siRNA treatment significantly increased the sprout length compared to the Bmp9 stimulation alone (Fig. 2D-E). An increasing trend was observed also for the number of sprouts per bead, such that no significant difference was present between the control samples (without BMP9) and the LFng siRNA samples with BMP9 stimulation. Taken together, these results strongly suggest that LFng mediates the expression of Notch target genes induced by Bmp9, which in turn regulates the inhibition of angiogenesis caused by Bmp9. The trends observed in the simulations (Fig. 2B) and the results obtained with the bead assay are strikingly consistent (Fig. 2C-D), strongly suggesting that the regulation of branch length and density is driven by changes in the Notch patterning temporal dynamics, allowing more tip cells to be selected faster (or slower) to generate more (or less) well extended sprouts (Fig. 2).

### Bmp9 and LFng are temporal regulators of tip/stalk cell shuffling

Endothelial cells continually rearrange positions during sprouting, with tip cells overtaken on average every 4-6 hours (Jakobsson *et al*., 2010; Arima *et al*., 2011). This process was shown to be Notch regulated, as Notch impacts VE-cadherin turnover; the Notch-mediated salt-and-pepper patterns create differential adhesions between individual ECs, contributing to their ability to rearrange positions (Bentley *et al*., 2014). Accordingly, we previously found that slower emergence of the Notch salt-and-pepper pattern had a knock-on effect and slowed cell overtaking rates in simulations and *in vitro* validation experiments (Kur *et al*., 2016). Thus, we next investigated whether Bmp9 could impact the dynamics of tip/stalk cell shuffling via LFng. To this aim, we adapted our previously developed and well validated agent-based model (the “memAgent-spring” MSM model) of cell shuffling (Bentley *et al*., 2014; Cruys *et al*., 2016; Kur *et al*., 2016). Similar to the ODE model utilized thus far, the MSM model captures Vegf/Notch signaling as per Fig. 1A, although using spatial agent-based modelling which captures cell shape, cell-cell adhesion and filopodia formation explicitly while using simple signaling rules rather than ODEs. This enables tracking of the position of cells as they rearrange positions within a sprout exposed to a gradient of Vegf along the sprout longitudinal direction (Fig 3A).

**Figure 3:**
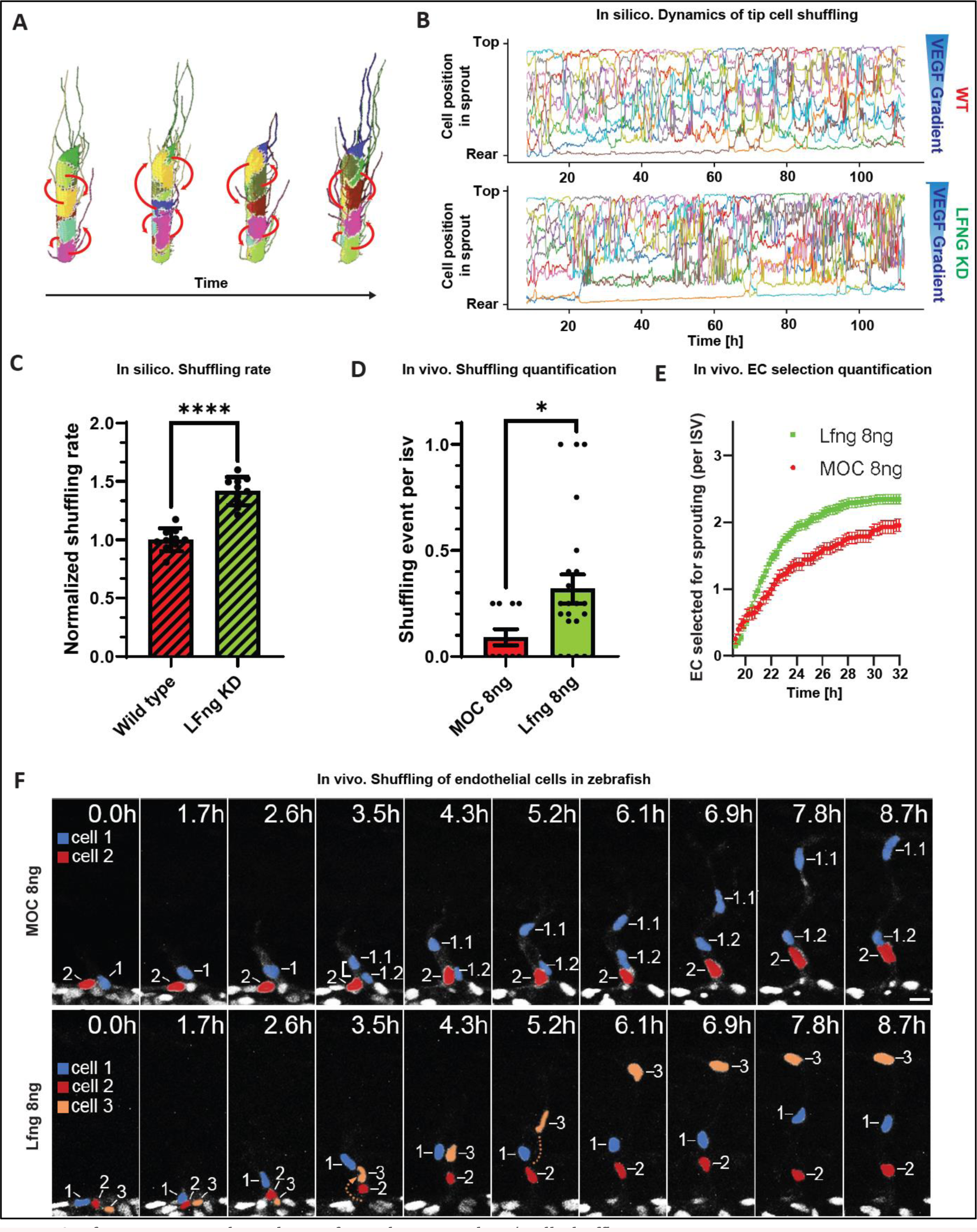
Lfng is a temporal regulator of tip selection and tip/stalk shuffling. (A) Representative simulation of the “memAgent-spring” MSM model. Over time (horizontal axis) cells on a sprout, each indicated with a different color, form filopodia (the lines extending from the sprout) towards the Vegf source (top) and compete for the top position of the sprout by exchanging positions (“shuffling”, red arrows). (B) Representative quantification of the coordinates of the center of mass of different cells in the memAgent-spring MSM model, indicated with different colors, with respect to the sprout longitudinal axis (vertical axis) over time (horizontal axis). Simulations were performed with (bottom graph) or without (top graph) Lfng KD. (C) Quantification of the shuffling rate resulting from the memAgent-spring MSM model with and without LFng KD, normalized over the control rate (wild type). The number of independent simulations per condition was N=10. (D-F) Quantification of EC nuclei shuffling (D), the number of ECs that are selected to branch (E), and representative time-lapse images of EC nuclei dynamics in intersegmental vessel (ISV) sprouts (F) in either control (MOC) or lfng knockdown (lfng) Tg(kdrl:nlsEGFP)^zf109^ embryos from 19 hours post-fertilization (hpf). In panel F, nuclei are pseudocolored according to the order of their emergence from the dorsal aorta (blue: first; red: second; yellow: third), with dividing cells retaining the same color and primary number as their parental cell (e.g. 1.1 and 1.2, top row). Bracket indicates a dividing cell. Shuffling events are indicated via dashed lines. Scale bar, 20 µm. Lfng morpholino-mediated knockdown increases the number of shuffling events (D, N = 11 for MOC, N = 23 for lfng), as well as the number of ECs selected for sprouting per ISV (E). All values are mean +/− SD. *p < 0.05; **p<0.01; ***p<0.001; ****p<0.0001; Mann-Whitney test.

LFng KD was modelled in the MSM by decreasing the likelihood of Dll4-Notch1 binding. The position of each cell along the sprout length was tracked for control and LFng KD conditions (Fig. 3B). Similar to previous computational results with Notch activity inhibition (Bentley *et al*., 2014), our simulations predicted that LFng KD strongly increased the frequency of tip/stalk shuffling (Fig. 3C). In particular, LFng KD weakened the feedback loops of the signaling network because of low Notch activation. As a result, cells at the tip of LFng KD sprouts could not efficiently inhibit their neighbors’ via Notch, allowing the neighbors to compete with and often overtake them, due to having higher Vegfr levels, motility, and adhesion turnover levels.

To verify these results, we performed *in vivo* experiments tracking EC nuclei dynamics during intersegmental vessel sprouting in zebrafish embryos (Fig. 3D-F, supplementary videos 1-2). Cell nuclei were tracked over time (Fig. 3F, supplementary videos 1-2) and, after the emergence of the first tip cell from the dorsal aorta (DA), the total number of shuffling events (Fig. 3D) and of emerging ECs (Fig. 3E) per intersegmental vessel (ISV) were quantified. Efficient KD of *lfng* in zebrafish was achieved using a previously validated morpholino oligonucleotide (MO) (Nikolaou *et al*., 2009). *Lfng* KD triggered significantly more cell shuffling than observed in embryos injected with a control MO, in confirmation of our computational results (Fig. 3C, 3D). Moreover, to define the number of ECs selected for sprouting, we quantified the number of ECs that emerged from the DA over the course of 13 h, excluding any additional cells in ISVs that arose from proliferation (Fig. 3E). A maximum of 2 emerging ECs per ISV were observed in control embryos, whereas a higher number of ECs emerged from the DA of *lfng* KD embryos (Fig. 3E), with some *lfng* KD ISVs exhibiting 3 emerging ECs (Fig. 3F). These observations are consistent with the concept that *lfng* KD decreases Notch activity leading to faster selection of more tip cells, as seen in our simulations (Fig. 2A, S2A, S2E). Representative examples of these results are shown in Fig. 3F. In the control case, a single tip cell (1, blue nucleus) emerges from the DA and, after cell division, is followed by the resulting cell (1.2, blue nucleus) and a stalk cell (2, red nucleus), without any shuffling (Fig. 3F, first row). In the case of *lfng* KD, three cells were selected to emerge, and the third cell (3, orange nucleus) sequentially overcame the first two cells thereby reaching the tip of the sprout (Fig. 3F, second row). Overall, these *in vivo* results demonstrate that LFng regulates the temporal dynamics of EC tip selection and shuffling.

### LFng loss can explain dysregulated cell rearrangement resulting from impaired Bmp9-Alk1 signaling

Next, we investigated whether the upregulation of LFng that is induced by Bmp9 could have a physiological relevance in the vascular dysfunctions associated with impaired Alk1 signaling. To this aim, we further investigated the role of LFng in the shuffling of ECs at the tip of sprouts, by simulating mosaic *in silico* experiments where cells were randomly assigned either a wildtype (red) or LFng KD (green) cell type. The dynamics of shuffling for mixed EC populations were compared to the shuffling dynamics obtained when only wildtype (green and red) cells are present in the sprout, as a control (Fig. 4A-B). As expected, consistent with the unbiased level of LFng in each cell, the control case presented random shuffling of cells (Fig. 4A, second row). On the other hand, in the mosaic simulations, LFng KD cells were prone to cluster at the tip of the sprouts, leaving the wildtype cells outcompeted at the far end of the sprout (Fig. 4A, first row). These results can be explained by the fact that the Notch receptors present in cells with LFng KD have a lower likelihood to bind to the Dll4 present in neighboring cells; therefore, they have a generally lower Notch activation compared to wildtype cells and their migratory behavior is thus less inhibited. A parameter sweep performed with a gradual increase in LFng KD resulted in a consistent, gradual clustering of mutated cells (Fig. 4B, in red). These computational results suggest that, during angiogenesis, cells with a decreased level of LFng have an advantage to localize and cluster at the sprout front.

**Figure 4:**
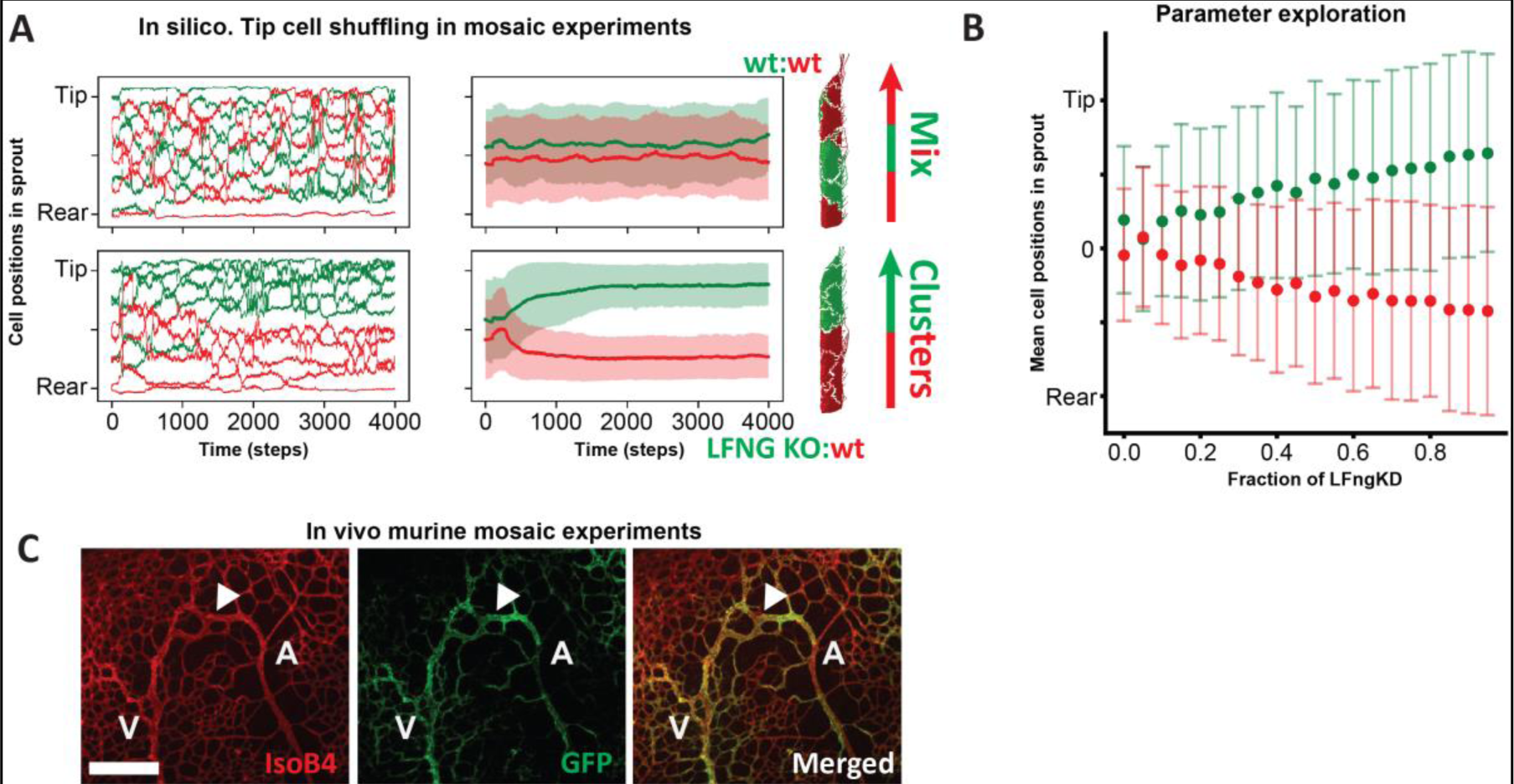
LFng KO cells spatially cluster in in silico and in vivo mosaic vessels and AVMs. (A) Representative (left) and mean +/− SD (right) positions of the center of mass of cells with respect to the sprout longitudinal direction (vertical axis) as computed by the memAgent-spring MSM model for mixed populations of cells, having Lfng KO (green, top row) or a wildtype phenotype. Over time (horizontal axis), LFng KO and wildtype cell cluster at the top and bottom of the sprout, respectively (top row). (B) Mean +/SD position (vertical axis) of the center of mass of cells in a sprout composed of a mixed population of cells (wildtype = red, LFng KD = green) as computed by the memAgent-spring MSM model. The fraction of LFng that was KD in each simulation group is indicated along the horizontal axis. (C) IsolectinB4 (red) and GFP (green) staining of wild type and Alk1Δ ECs, respectively, in retinal vessel mosaic experiments. The arrowhead points to the AVM, which is almost exclusively composed of Alk1 KO/GFP-positive cells, as opposed to the rest of the vascular plexus where there is a high degree of heterogeneity. Scale bar, 500 µm. All values are mean +/− SD.

We have previously shown that, in mosaic sprouts with control and Alk1 knockout (KO) cells, the Alk1 KO cells preferentially localize at the tip of the sprout (Larrivée *et al*., 2012). While we have previously ascribed this clustering to the direct targeting of Hey1 by Alk1 (Larrivée *et al*., 2012), our present simulations indicate the loss of LFng in Alk1 KO cells as another contributing mechanism. This mechanism might also be relevant for disease settings. In *in vivo* mosaic experiments (Fig. 4C), we observed that GFP Alk1 KO cells formed arteriovenous malformations (AVMs), consistent with previous studies showing that Alk1 dysfunction is involved in these malformations (Baeyens *et al*., 2016), which tend to be comprised of cells low in Notch signaling (Tual-Chalot *et al*., 2014). AVMs are characteristics of HHT, which can indeed be phenotypically copied by Alk1 KD (Arthur and Roman, 2022). These results point therefore to a possible relevant role of LFng in the context of HHT.

### Bmp9-mediated LFng and Jag1 expressions in ECs ensure contemporary Notch activation in co-cultured ECs and pericytes

Bmp9 is known to strongly upregulate the expression of Jag1 (Morikawa *et al*., 2011), another ligand of the Notch pathway that is highly involved in cardiovascular (and angiogenesis) regulation (Loomes *et al*., 1999; High *et al*., 2008; Bauer, Jackson and Jiang, 2009; Benedito *et al*., 2009; Pedrosa *et al*., 2015). Previous research has shown that LFng-mediated Notch1 glycosylation inhibits Jag1-Notch1 activation (Yang *et al*., 2005; Kakuda and Haltiwanger, 2017; Kakuda *et al*., 2020), and the expression of Jag1 is pro-angiogenic (Benedito *et al*., 2009). The contemporary increase of Jag1 and LFng that is induced by Bmp9, together with the anti-angiogenic role of Bmp9, thus appear counterintuitive. Jag1 plays a crucial role for the signaling crosstalk of ECs with pericytes (Liu, Kennard and Lilly, 2009; Tefft *et al*., 2022), fundamental for endothelial stabilization (Geevarghese and Herman, 2014; Dibble *et al*., 2023). Therefore, to reconcile Bmp9-mediated LFng and Jag1 upregulation, we investigated whether LFng plays a role in this context. Specifically, we hypothesized that Jag1 and LFng act in synergy to potentiate the crosstalk between ECs and pericytes, where LFng acts as railroad switch diverting Jag1 from activating Notch1 on ECs, to activate Notch3 on pericytes, allowing the high Jag1 levels to improve, rather than hinder, vessel stability.

Motivated by previous experiments (Benedito *et al*., 2009; Larrivée *et al*., 2012; Kakuda and Haltiwanger, 2017; Kakuda *et al*., 2020; Tefft *et al*., 2022), we extended our ODE model of EC signaling by adding the EC crosstalk with pericytes as mediated by Jag1 (Fig. 5A). In the model, Bmp9 upregulated EC expression of Jag1 (Larrivée *et al*., 2012), which could in turn bind and activate Notch1 in ECs and Notch3 in neighboring pericytes (Benedito *et al*., 2009; Tefft *et al*., 2022). Notch3 activation induced pericytes to express Dll4 (Tefft *et al*., 2022), which in turn could activate Notch1 in ECs establishing a full feedback loop between pericytes and ECs. Based on previous experiments (Taylor *et al*., 2014; Kakuda and Haltiwanger, 2017; Kakuda *et al*., 2020), and differently than previous computational models (Boareto, Jolly, Ben-Jacob, *et al*., 2015; Boareto, Jolly, Lu, *et al*., 2015), LFng was assumed to increase the binding of both Dll4 and Jag1 to Notch1, while decreasing the chance of Jag1-Notch1 activation post-binding (Fig. 5A).

**Figure 5:**
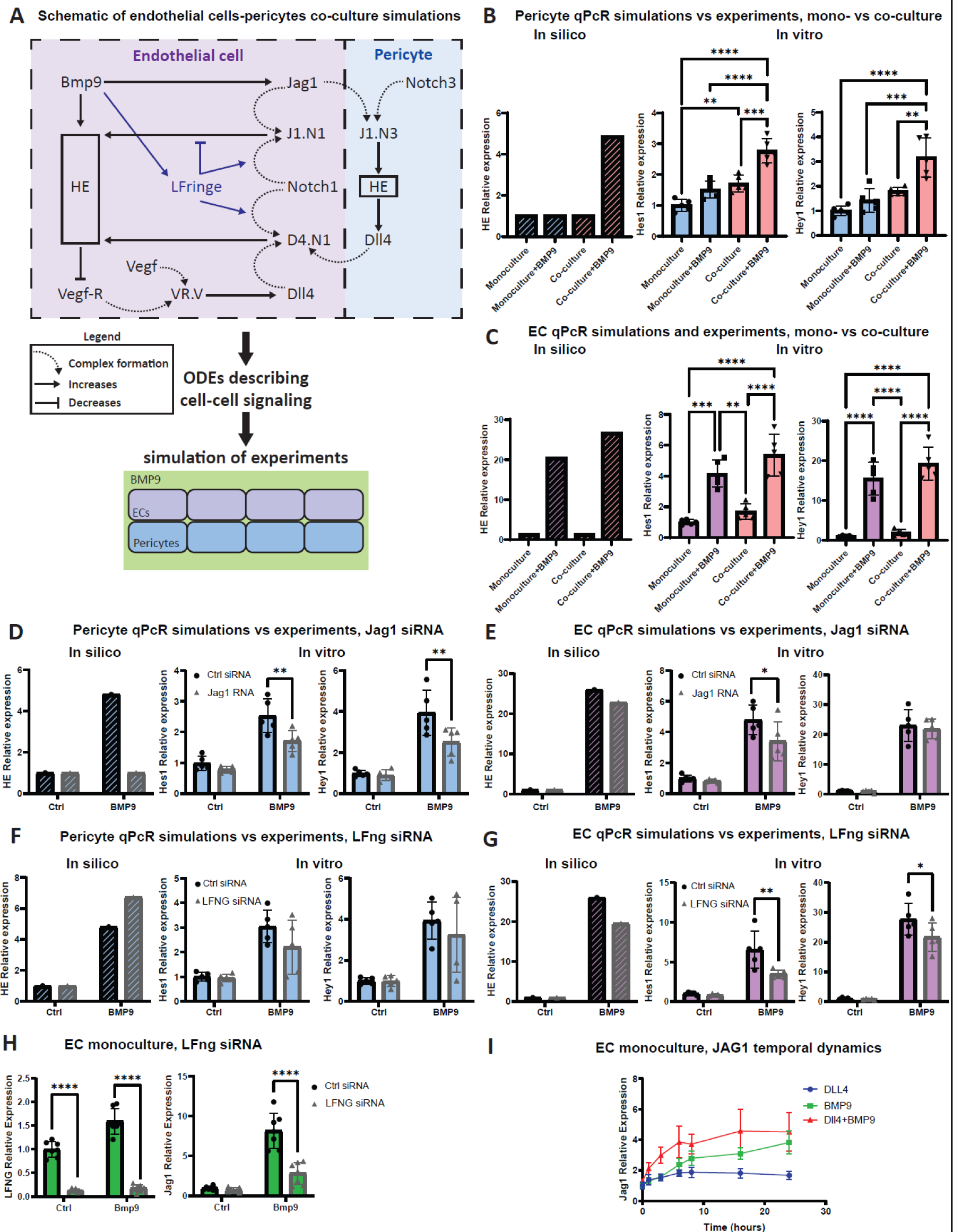
Bmp9-mediated LFng upregulation potentiates EC-pericyte interactions. (A) Schematic representation of the signaling crosstalk between ECs and pericytes, as well as of the simulation and in vitro experiments. Full names correspond to unbound ligands and receptors (e.g. Notch1), while capital letters with numbers correspond to bound receptors (e.g. N1.D4 is the bound Notch1-Dll4 complex). Straight and curved arrows represent upregulation and protein complex formation, respectively, while the ⊣ symbol represents inhibition. (B-G) Experiments (full color) and simulations (striped color) of qPCR analysis of Hes1 and Hey1 in HUVECs and pericytes, cultured alone or in co-culture, induced by Bmp9 injection (10 ng/ml) after 24h stimulation, with (bright color) or without (shaded color) Jag1 or LFng siRNA treatment. The number of independent experimental repeats was N = 5. (H) qPcR analysis of LFng and Jag1 in HUVECs induced by Bmp9 injection (10 ng/ml) after 24h stimulation, with or without LFng siRNA treatment. The number of independent experimental repeats was N = 7. (I) qPCR analysis of Jag1 in HUVECs stimulated by Bmp9 (10 ng/ml) and/or Dll4 (10 ng/ml) at different time points, up to 24h. The number of independent experimental repeats was N=7. All values are mean +/− SD. *p < 0.05; **p<0.01; ***p<0.001; ****p<0.0001; ANOVA.

To investigate the possibly synergistic roles of LFng and Jag1, we simulated and performed *in vitro*, mono- and co-culture qPCR experiments of HUVECs and pericytes, stimulated with Bmp9, with and without downregulation of LFng or Jag1 in ECs. In agreement with our experimental findings, the simulations indicated that Bmp9 has little to no effects on Notch activation in pericytes unless co-cultured with ECs, for which a strong increase in Hey1/Hes1 was predicted (Fig. 5B). In the simulations, co-culture also increased the previously observed Hes1/Hey1 upregulation in ECs induced by Bmp9. This was not observed in the experiments, although a non-significant increasing trend was present (Fig. 5C). Overall, the experiments and simulations showed that BMP9 increases Notch signaling activation in pericytes through its effects on ECs.

In the simulations, Bmp9 effects are entirely mediated by Jag1 present in ECs. Accordingly, knocking down Jag1 in ECs was predicted to inhibit BMP9 effects on pericytes in co-culture conditions (Fig. 5D), and some decreasing effects were predicted also for ECs (Fig. 5E). This was confirmed by qPCR experiments with partial KD of Jag1, which caused a decrease of Hes1 and Hey1 in pericytes (Fig. 5D), and of Hes1 in ECs (Fig. 5E). Finally, to validate our hypothesis of the importance of LFng in this context, we performed LFng KD experiments and simulations. In the model, as a result of LFng KD, Jag1 had a decreased binding rate to Notch1 in ECs. Therefore, more Jag1 was available for Notch3 binding and activation in pericytes, which exhibited increased levels of Hes1/Hey1 (Fig. 5F). In ECs, given that Dll4 as well had decreased Notch1 affinity, LFng KD caused a decrease in Hes1 and Hey1 expression. The experiments were in partial agreement with the simulations, showing that LFng KD in ECs decreases Hes1 and Hey1 expression in ECs as a result of Bmp9 (Fig. 5F), in agreement with our previous results (Fig. 1). However, it does not have significant effects on pericytes (Fig. 5E); actually, a non-significant downregulating trend can be observed.

This disagreement suggested the existence of other underlying mechanisms. Given the important role of EC Jag1 for pericytes, we checked whether LFng KD affects Jag1 expression in ECs. Interestingly, we observed that LFng mediates the Bmp9-induced upregulation of Jag1 (Fig. 5H); in fact, LFng KD strongly decreased Jag1 expression resulting from Bmp9. This explained the apparent disagreement between experiments and simulations present in Fig. 5E. Specifically, in the model, as a result of LFng KD, Jag1 had a decreased binding rate to Notch1 in ECs; therefore, more Jag1 was available for Notch3 binding and activation in pericytes, which exhibited increased levels of Hes1/Hey1 (Fig. 5F). In the experiments, more Jag1 was not available for Notch3 binding because of a decreased Jag1 expression in ECs, resulting from LFng KD, such that Hes1/Hey1 expression was unaffected (Fig. 5E). The role of LFng in regulating Jag1 expression in response to Bmp9 suggests that a synergy between Bmp9 and Notch signaling might exist also in this context. Indeed, a time course qPCR analysis of Jag1 expression in HUVECs confirmed that Dll4 coating can induce a moderate increase of Jag1 *in vitro* and, when combined with Bmp9 stimulation, this leads to a stronger increase compared to Bmp9 alone (Fig. 5I). Overall, by combining experiments and simulations, we showed that Bmp9 upregulation of LFng ensures high Dll4-Notch activation in ECs despite also leading to increased EC Jag1 expression, which is fundamental for the Bmp9- and EC-mediated Notch activation in pericytes.

### LFng and Jag1 induced by Bmp9 act in synergy to stabilize the vasculature

Pericytes are fundamental for the stabilization of ECs and associated vasculature (Geevarghese and Herman, 2014; Lebrin, 2015; Dibble *et al*., 2023). Therefore, we next computationally investigated whether LFng and Jag1 might act in synergy to stabilize ECs via pericyte interaction. Given the importance of LFng for the temporal regulation of tip-stalk cell selection, we hypothesized that both LFng and Jag1 play a role in the temporal regulation of EC stabilization. To test this hypothesis, we performed simulations where a row of ECs was first exposed to VEGF for 24h, to establish a stable tip/stalk pattern, followed by 24h of exposure of VEGF and Bmp9, with and without LFng and/or Jag1 inhibition, with the addition of pericytes (Fig. 6A). These simulations were adopted to approximate the final stages of angiogenesis, where ECs recruit pericytes and start being exposed to higher levels of Bmp9 present in the blood (David *et al*., 2008). To better track the different roles of the two proteins, the parameters of Jag1 and LFng were still modelled independently as in the original model, despite LFng involvement in Jag1 regulation (Fig. 5H). In the simulations, vessels were assumed to be stable when all ECs exhibited a stalk phenotype. Without Bmp9, despite the addition of pericytes, ECs retained the previously established tip-stalk pattern, without vessel stabilization. Bmp9 addition caused an increase in LFng and Jag1 expression, with a consequential increase of Hes1/Hey1, in agreement with our previous findings (Fig. 5). This was followed by Vegfr2 and filopodia inhibition and, therefore, by all ECs stabilizing by exhibiting a stalk phenotype (Fig. 5D). In agreement with our hypothesis, the simulations showed that LFng and Jag1 expressions resulting from Bmp9 positively correlated with the rate of stabilization (Fig. 6B-D). Both proteins were essential for this process; decreasing either LFng or Jag1 below a threshold led to a loss of stabilization, with ECs retaining the tip-stalk pattern (Fig. 6B, bright green area). Therefore, our simulations strongly suggest that LFng and Jag1 act in synergy to potentiate the Notch crosstalk between ECs and pericytes, to upregulate Notch activity in EC and drive the stabilization of the vasculature.

**Figure 6:**
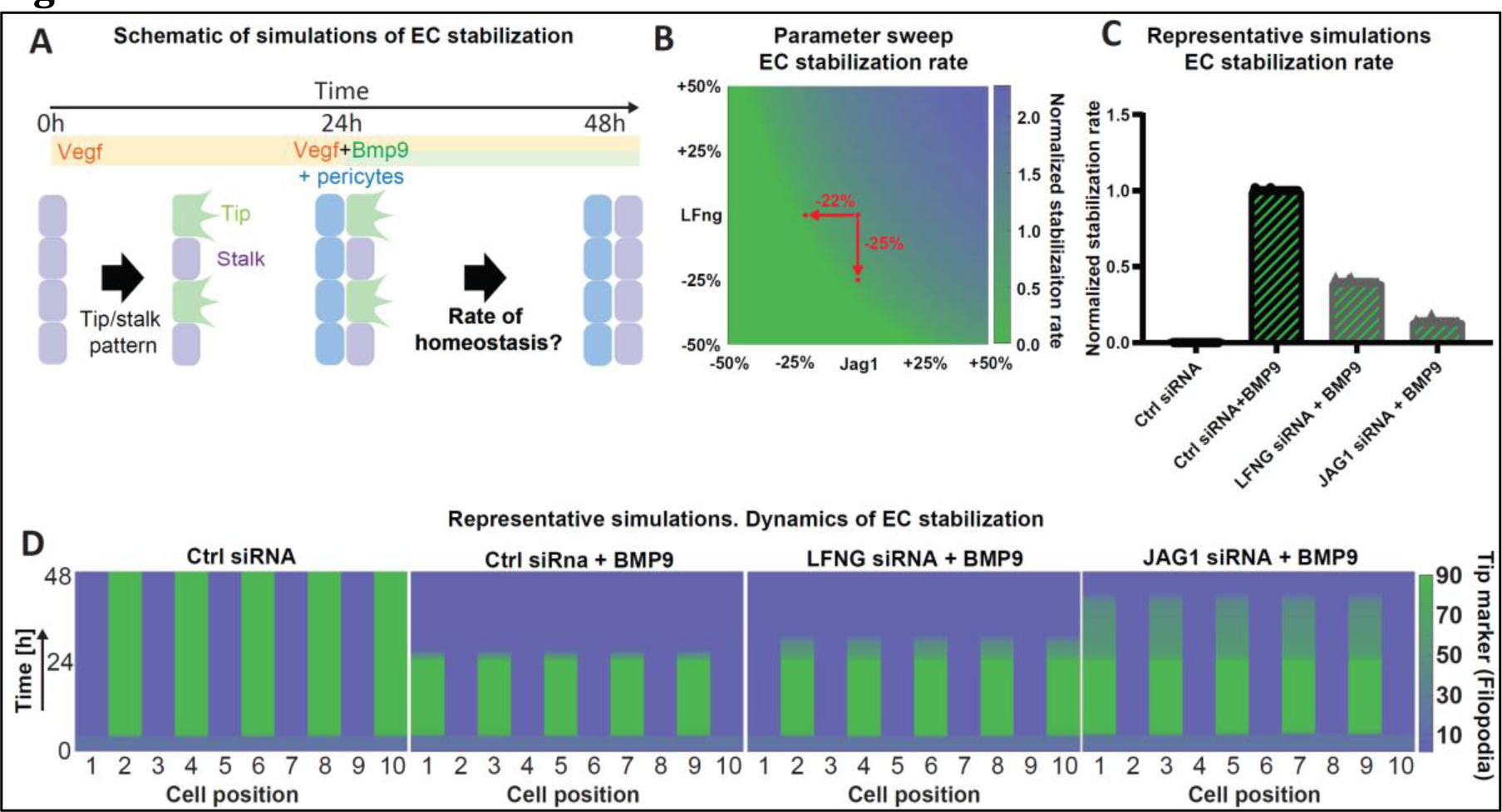
LFng and Jag1 as temporal regulators of endothelial stabilization. (A) Schematic representation of the computational simulations performed to compute the rate of EC stabilization upon exposure of Bmp9 and pericyte contact, 24h after Vegf exposure. (B) Average normalized rate of EC stabilization for varying values of Bmp9 effects on LFng (vertical axis) and Jag1 (horizontal axis) with respect to original parameters (central red dot). Two additional dots were placed corresponding to the representative LFng and Jag1 siRNA simulations reported in (C), with the associated parameter downregulations indicated with arrows. The color scheme represents the normalized rate of EC stabilization. (C-D) Representative simulations of EC stabilization for a row of 10 ECs exposed to Vegf for 24h and, for the subsequent 24h, Bmp9 and pericyte contact. In these representative simulations, LFng and Jag1 siRNA were mimicked by reducing the associated parameters 25% and 22%, respectively. The color scheme in (D) represents the amount of filopodia in each cell. The EC stabilization rate in (C) was approximated as the ratio between the time necessary for EC stabilization in the simulation in exam, normalized over the time necessary for the original parameters. The number of simulations for each condition was N = 50.

## Discussion

Bmp9-Alk1 and Notch signaling are interconnected in multiple cellular systems (Mostafa *et al*., 2019). By integrating experiments with computational simulations, we here provide a key missing link that Bmp9 regulates LFng, which enables a positive feedback multiplicative crosstalk between Bmp9 and Notch signaling than better explains observations than the prevailing independent, additive model of bmp9-Alk-Notch crosstalk (Larrivée *et al*., 2012). Furthermore we find that the upregulation of LFng contributes to: 1) the regulation of the temporal dynamics of angiogenesis exerted by Bmp9 over the short timeframe of hours, impacting branch length/density, cellularity and spacing; 2) the non-physiological EC rearrangement at the tip of sprouts in Alk1 loss-of-function conditions, indicating a role in pathological AVM formation in Alk1 mutants; 3) the upregulation of Jag1 and diversion of Jag1 to bind Notch3 on pericytes rather than Notch1 on ECs, contributing to vessel stabilization and providing a new explanation for decreased pericyte coverage and EC Notch activation observed in Alk1 mutant AVMs.

LFng has been shown to play crucial roles for several developmental and cellular processes (Cole *et al*., 2002; Morales, Yasuda and Ish-Horowicz, 2002; Dale *et al*., 2003; Serth *et al*., 2003; Tsukumo *et al*., 2006; Shifley *et al*., 2008; Kato *et al*., 2010; Xu *et al*., 2010; Yuan *et al*., 2011; Semerci *et al*., 2017; Derada Troletti *et al*., 2018) due to its dominant effect on Notch regulation (Pennarubia *et al*., 2021). Temporally cyclic regulation of LFng expression has been reported to regulate the oscillatory nature of Notch signaling during mesodermal segmentation (Cole *et al*., 2002; Morales, Yasuda and Ish-Horowicz, 2002; Dale *et al*., 2003; Serth *et al*., 2003; Shifley *et al*., 2008). In the context of angiogenesis, we have previously shown that the Notch temporal dynamics influences endothelial cell fate selection and shuffling, and the resulting vascular topology (Bentley and Chakravartula, 2017). Several temporal modulators of Notch signaling have been identified, including Dlk1 and Mash1 (Yun *et al*., 2002; Finn *et al*., 2019). During angiogenesis, LFng inhibition leads to increased tip-cell selection and vascular density (Benedito *et al*., 2009). Combined with our previous simulations (Venkatraman, Regan and Bentley, 2016), our new experimental and computational results strongly indicate that LFng influences angiogenesis by acting as a temporal modulator of endothelial cell fate selection and rearrangement (Figs. 2-3). These mechanisms might as well explain some of the effects that Bmp9 has on angiogenesis, as demonstrated by the partial rescue of Bmp9-stimulated angiogenic sprouts by LFng KD (Fig. 2). Therefore, while the cyclic activity of Bmp9 over the course of days has been shown to be central for the regulation of angiogenesis (Guihard *et al*., 2020), our study also points at Bmp9 as a regulator of the temporal dynamics of angiogenesis in the shorter timescales of hours.

It is well recognized that the regulation of angiogenesis and vasculature homeostasis exerted by Bmp9-Alk1 signaling is central in health and disease. Dysregulation of Alk1 can lead to AVMs, characteristics of HHT (Larrivée *et al*., 2012; Tual-Chalot *et al*., 2014). Via mosaic experiments, we have previously shown that Alk1 KO cells cluster at the tip of sprouts (Larrivée *et al*., 2012), and that AVMs in mosaic *in vivo* experiments are mainly formed by the Alk1 KD cells (Baeyens *et al*., 2016). Our simulations indicate that the latter phenomenon can be explained by the downregulated expression of LFng in Alk1 KO cells (Fig. 4), which makes them less prone to be inhibited by Dll4 present in neighboring wild type cells. Notch inhibition can cause AVMs (Krebs *et al*., 2004; Nielsen *et al*., 2014). AVM regions as a result of Alk1 KD are typically characterized by lower Notch activity compared to physiological values (Tual-Chalot *et al*., 2014), which could also be caused by a downregulation in LFng. This downregulation might not only impair Notch among ECs, but also between ECs and pericytes, as demonstrated by our co-culture results (Fig. 5). Interestingly, AVM patients exhibit lower pericyte coverage around the vessels (Winkler *et al*., 2018). Moreover, poor Notch signaling in pericytes has been shown to lead to AVM formation (Nadeem *et al*., 2020). Our co-culture data strongly suggests that the Bmp9-mediated upregulation of Jag1 is fundamental for Notch activation in pericytes, and that LFng contributes to this Jag1 upregulation by increasing Notch activity in ECs (Fig. 5). These data are also in accordance with a previous study showing that endothelial-expressed Jag1 is required for Notch3 signaling in perivascular cells (Liu, Kennard and Lilly, 2009). Overall, our data thus point at Bmp9 as a potentiator of the EC-pericyte Notch signaling communication, via upregulation of both Jag1 and LFng, which act in synergy to induce elevated Notch activation in both pericytes and ECs, thereby facilitating cellular homeostasis and vessel stabilization (Fig. 6). These mechanisms are likely to be key for vasculature homeostasis and, when disrupted, might be linked to AVMs and HHT.

Smad1/5 have been previously identified as regulators of LFng in ECs (Morikawa *et al*., 2011). Mechanistically, therefore, it is likely that Bmp9 activates Alk1, whose signal is transmitted via Smad1/5, with a consequential upregulation of LFng (Fig. 1A). Similar effects observed for ERG, previously shown to upregulate LFng in HUVECs (Shah *et al*., 2017), could also be explained with analogous mechanisms, given the promotion of Smad1 transcriptional activity consequential to ERG (Dufton *et al*., 2017). In contrast, we showed that Dll4-mediated Notch activation did not induce relevant LFng expression changes in HUVECs (Fig. 1), differently than in somites (Del Barco Barrantes *et al*., 1999; Morales, Yasuda and Ish-Horowicz, 2002), thereby highlighting the key role of Bmp9-Alk1 signaling for LFng regulation in ECs.

Taken together, these data uncover a novel role for BMP9-mediated LFng expression in maximizing Notch1 activation by Dll4 in endothelial cells, while concomitantly maintaining Notch3 signaling in pericytes through Jag1, overall demonstrating the functional relevance of the Bmp9-mediated LFng upregulation at different timescales of the angiogenic process in health and disease.

## Methods

### Experimental methods

#### Antibodies and recombinant proteins

Antibodies against LFng (CST# 66472), Cleaved Notch1 (NICD CST# 4147), Jag1 (AF599) and β-actin were purchased from Cell Signaling, R&D systems and Santa Cruz Technologies respectively. Quantitect qPCR primers for Jag1 (QT00031948), Hey1 (QT00035644), Hey2 (QT00026971), Dll4 (QT00081004), Hes1 (QT00039648), LFng (QT00212240), MFng (QT00084007), and RFng (QT01160075) were purchased from Qiagen, Recombinant BMP9 (3209-BP-010) and Dll4 (1506-D4-050) were obtained from R&D systems.

#### Cell culture

HUVECs from Promocell (pooled donors, Cat#C-12203) were cultured with ECGM-2 (Lonza). Before stimulation with BMP9 10 ng/ml, HUVECs were starved overnight in EBM-2 (Lonza) supplemented with 0.1% FBS. The same was done before stimulation with sDll4, which was precoated on 6-well plates at 10 µg/ml as previously described (Larrivée *et al*., 2012).

#### siRNA transfection

For LFng KD *in vitro*, HUVECs were transfected with siRNA obtained from QIAGEN (Hs_LFNG_13 FlexiTube siRNA; Cat# SI05348756), using the RNAiMax reagent (ThermoFisher, catalog# 13778030) according to the manufacturer’s instructions, 24 hours prior to stimulation with BMP9. The same procedure was used for the Jag1 KD (Hs_JAG1_5 Flexitube siRNA Cat#: SI02780134). Negative control experiments were performed with AllStars Negative Control siRNA from QUIAGEN (Cat# 1027280).

#### Pericyte co-cultures

250,000 HUVECs previously transfected with control siRNA, or siRNA targeting JAG1 or LFNG, were grown in ECGM-2 medium supplemented with VEGF for 24 hours before the addition of 100,000 human pericytes (Promocell, Cat# C-12980), with or without BMP9. After 24 hours, endothelial cells were separated from pericytes using CD31-labeled Dynabeads, and mRNA was harvested using RNeasy MiniKit (QIAGEN). Relative purity of HUVECs and pericytes was confirmed by qPCR analysis of endothelial (PECAM1, CDH5) and pericyte (PDGFRB) markers. As controls, 100,000 pericytes were cultured with or without BMP9 in the absence of HUVECs.

#### Fibrin gel bead assay

The bead assay was performed following previous protocols (Nakatsu and Hughes, 2008; Newman *et al*., 2011). Briefly, Cytodex microcarrier beads were purchased from (Sigma-Aldrich, St. Louis, MO). Prior to use, the beads were swollen in PBS (50 mL/g) and autoclaved at 120°C, to obtain a solution of 60,000 beads/ml. Beads were then coated with approximately 400 HUVECs per bead, and transferred to a T25 flask in 5mL of EGM-2. After a day, a (2.5 mg/mL) fibrinogen solution in EBM-2 was prepared, adding aprotinin (Sigma-Aldrich) for a final concentration of 50 µg/ml. The beads were then seeded, for a final concentration of 500 beads/ml. Thereafter, 0.625 Units/mL of thrombin (Sigma-Aldrich) were added to the mix. Human Dermal Fibroblasts (HDF; Promocell, Cat# C-12302) were finally added on top of the fibrin gels, at a concentration of 20000 cells per well. The medium, supplemented with or without 10 ng/ml VEGF and/or 1 ng/ml BMP9 was replaced every 2 days, and sprouts were imaged by fluorescence after 7 days. Single sprouts were defined as continuous cellular protrusions departing from the beads that were longer than the bead diameter.

#### Zebrafish

Embryos and adults were maintained under standard laboratory conditions as described previously (Zakirov *et al*., 2021) and experiments were approved by the University of Manchester Ethical Review Board and performed according to UK Home Office regulations. The *Tg(kdrl:nlsEGFP)^s896^* strain was established previously (Blum *et al*., 2008). To knock down gene expression, embryos were injected at the one-cell stage with either 8 ng control MO (MOC) or 8 ng *lfng* MO (*lfng*). MO sequences were: 5’-CCTCTTACCTCAGTTACAATTTATA-3’ (MOC), 5’-ACCGTGTATACCTGTCGCATGTTTC - 3’ (*lfng*) (Nikolaou *et al*., 2009). All MOs were purchased from Gene Tools. Movies of zebrafish ECs were acquired as previously described (Costa *et al*., 2016). Briefly, live embryos were mounted in 1% low-melting agarose (containing 0.1% tricaine) in glass-bottom dishes and were continually perfused with embryo water supplemented with 0.0045% 1-phenyl-2-thiourea and 0.1% tricaine at 28 °C using an in-line solution heater and heated stage controlled by a dual-channel heater controller (Warner Instruments). Embryos were imaged using 40×-dipping objectives on a Zeiss LSM 700 confocal microscope.

### Mice

The Maisonneuve-Rosemont Hospital ethics committee, overseen by the Canadian Council for Animal Protection, approved all experimental procedures (protocol number: 2014-18). All the animal experiments were conducted according to the Standard Operation Procedures (SOP) of the Maisonneuve-Rosemont Hospital Animal Ethics Committee. Tamoxifen-inducible Cdh5-CreErt2 and acvrl1^loxP^ mice were kindly provided by Ralf Adams and S. Paul Oh respectively. To generate Alk1ΔEC mice, Cdh5-CreErt2 and acvrl1^loxP^ mice were crossed and injected with 50 mg/kg tamoxifen dissolved in corn oil at P4 and euthanized at P5. Retinas were harvested and stained with IsolectinB4 and selected antibodies as previously described (Larrivée *et al*., 2012). Throughout the studies, Cdh5-CreErt2-Alk1+/+ (thereafter referred as C5Cre) mice injected with tamoxifen as described above were used as controls.

### Isolation of Retinal Endothelial Cells

Following tamoxifen injections at P4, retinas from C5Cre-Rosa^mTmG^ and Alk1ΔEC-Rosa^mTmG^ were harvested at P5 and dissociated for 20 min using collagenase type II as previously described (Chavkin, Walsh and Hirschi, 2021). GFP-positive cells were harvested by FACS. Overall, 10,000 retinal endothelial cells were obtained from 4 pooled retinas. Sorted endothelial cells were resuspended in RNA lysis buffer and RNA was isolated (RNeasy MicroKit; QIAGEN) and processed for quantitative RT-PCR.

### Statistical analysis

All values are expressed as the mean ± standard deviation (SD). Statistical analyses were performed using the GraphPad Prism software. Quantitative differences between multiple groups were compared by using one-way ANOVA test, once normality and homogeneity was proved with Shapiro-Wilk test. A level of P <0.05 was considered statistically significant.

### Computational methods

#### ODE model of EC signaling

##### Model assumptions

The effects of BMP9 on the Notch-mediated interaction between ECs was simulated by extending a previous ODE model (Venkatraman, Regan and Bentley, 2016). The original model was developed based on mass-action kinetics equations. Briefly, the original model assumes that Notch can bind and be activated by Dll4 present in adjacent cells. Notch activation leads to upregulation of Hey1, which in turn downregulates the expression of VEGFR2. VEGFR2 can bind and be activated by VEGF present in the environment, which leads to upregulation of Dll4 and filopodia formation. Finally, filopodia formation is assumed to increase the VEGF that is detected by the cell and thus available for VEGFR2 binding and activation. Here, this original model was extended by accounting for the possible movement of Notch proteins on the cell membrane, to enable the simulation of a row of ECs. Similar to previous studies (Koon *et al*., 2018; Fisher and Strutt, 2019), the cell membrane was considered as divided between two edges, left and right, while the movement of unbound Notch proteins between the two edges was included by assuming random movement, and thus by adding an equation term similar to diffusion. For example, for the content of a protein M, the term Δ_*L*_ defines the variation over time of the protein content *M*_*L*_ on the left edge as:

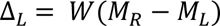

where *W* is a coefficient scaling the movement rate of the proteins between the two edges. An analogous expression was adopted for Δ_*R*_ on the right edge.

The upregulation of Bmp9 on Hes1/Hey1 expression and the Notch receptor-ligand binding rate was modeled linearly as follows:

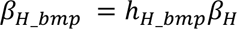

and

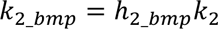

where *β*_*H*_*bmp*_ and *k*_2_*bmp*_ are respectively the Hes1/Hey1 expression and the Notch receptor-ligand binding rate for varying values of Bmp9 concentration; *β*_*H*_ and *k*_2_ indicate the corresponding homeostatic values; while ℎ_*H*_*bmp*_ and ℎ_2_*bmp*_ are parameters that scale the influence of Bmp9 on these two biological phenomena. The values ℎ_*H*_*bmp*_ = 1 and ℎ_2_*bmp*_=1 were chosen to simulate physiological conditions. Values larger than 1 were chosen to simulate Bmp9 injection or Alk1 overexpression. Values lower than 1 were chosen to capture the effects of Alk1 KD. All the other model parameter values were kept as in previous studies (Venkatraman, Regan and Bentley, 2016), unless specified.

The model was coded in Matlab2019b and solved using ODE15s. The ODEs are deterministic, but the final solutions depend on the initial conditions because of the multiple attractors present in the solution space and the limited simulation time. To account for this, similar to previous studies with analogous Notch models (Boareto, Jolly, Lu, *et al*., 2015; Loerakker *et al*., 2018; Ristori *et al*., 2020; Tiemeijer *et al*., 2022), the simulations were performed for 10000 random initial conditions. Simulations were performed for 10 adjacent cells with periodic boundary conditions, as in a recent study (Tiemeijer *et al*., 2022), such that the first cell was assumed to be interacting with the last (10^th^) cell. For more details on the specific ODEs, we refer the reader to the Supplementary Information.

##### Simulations

*In vitro* qPCR experiments with EC monolayers were simulated by assuming that cells were not exposed to VEGF (*V* = 0). In agreement with the actual experiments, the ODEs were solved with a final time equal to 48h, and external Dll4 and/or BMP9 were only added 24h hours after the start of the experiment. The parameter representing external Dll4 (*D*_*ext*_) was calibrated such that the Hey1 expression predicted for Dll4-stimulated cells is 5 times higher than the Hey1 expression predicted for starved cells, in agreement with previous *in vitro* results (Larrivée *et al*., 2012). The parameter sweep adopted to explore the possible effects of BMP9 on Notch was performed by multiplying and dividing by two the model parameters associated with Notch signaling (e.g. Notch-Dll4 binding rate, Notch production, etc.). For the specific parameters that were varied during the parameter sweep, we refer the reader to Eqs. S1-S5 and Fig. S1. After the identification of the effects of Bmp9 on the Notch receptor-ligand binding rate as a possible candidate for explaining previous qPCR experiments (Larrivée *et al*., 2012), ℎ_*H*_*bmp*_ and ℎ_2_*bmp*_ were calibrated to mimic such experiments. The other parameters were left unchanged.

Tip/stalk selection at the onset of angiogenesis was simulated by adopting the same ODE model. Cells were exposed to no VEGF for the first 24h, thereby mimicking homeostatic conditions, while cells were exposed to relatively high VEGF levels (*V*_*ext*_ = 0.05 units) for the subsequent 24h. For Fig. 2A-B, motivated by our experiments and simulations, Bmp9 injection was modeled by increasing the basal expression of Hes1/Hey1 and LFng by four times (ℎ_*H*_*bmp*_ = 4, and ℎ_2_*bmp*=_ = 4), while LFng KD was modeled by dividing the control value by 1.5 (ℎ_2_*bmp*_ = 1/1.5).

##### Analysis of in silico tip-stalk pattern formation rate

For the *in silico* simulations, the time necessary to form the tip-stalk pattern that is characteristic of angiogenesis was evaluated by assigning fates to each cells based on their specific filopodia values, to be consistent with herein experiments and previous computational analysis (Bentley, Gerhardt and Bates, 2008). Endothelial cells were assigned a tip cell phenotype if their filopodia exceeded a certain heuristically-determined threshold (*f*_*min*_ = 25 *c*. *u*.), and a stalk cell phenotype otherwise. A tip-stalk pattern was assumed to be established when the percentage of tip cells was between 40% and 50%, and no adjacent tip cells were present. To obtain a proxy of the tip-stalk pattern formation rate, the time to pattern was normalized over 1 day, taken as a reference value.

### memAgent spring model (MSM)

Shuffling of cells at the tip of sprouts was simulated with the MSM (Bentley, Gerhardt and Bates, 2008; Bentley *et al*., 2009), incorporating Cellular Potts Model based modelling of cell rearrangement via differential adhesion (Bentley *et al*., 2014). This model and its parameters have been previously validated against several *in vivo* experiments in mouse and zebrafish (Bentley *et al*., 2014; Kur *et al*., 2016; Page *et al*., 2018). The model accounts for the movement of the cell membrane as influenced by VEGF, Notch signaling, filopodia formation and neighboring cells. Individual computational agents (memAgents) represent the EC membrane, which is held together by the actin cortex represented by springs following the Hooke’s law. Cell membrane movement, Dll4 content, and filopodia formation are proportional to the VEGF detected by the cell, which in turn decreases in response to Dll4-mediated Notch activation. In the present study, LFng KD was accounted for by scaling the model parameter representing Notch activation. In particular, to simulate reduced Bmp9 signaling in Alk1 mutant cells, we reduced Notch activation in mutant cells to 25% of its WT value.

In the simulations, the vessel was considered to be oriented in the direction of a VEGF gradient that was constant over time, so that excited tip-like cells moved along the vessel up the gradient. A vessel was composed of 10 cells competing for the tip position, initialized next to each other (one per vessel cross section). We allowed the vessel delta-notch pattern to stabilize for around 8 hours (1000 30 s timesteps) before allowing cells to rearrange via differential adhesion. We then continued the simulation for either 4000 timesteps (around 33 hours) to evaluate tip cell competition and 14000 timesteps (around 109 hours) to evaluate cell rearrangement. Model parameters were kept as in previous studies (Bentley *et al*., 2014; Kur *et al*., 2016), unless otherwise specified. As the model is stochastic, we quantified the average position of the center of mass along the VEGF gradient between mutant and wild type cells for 10 repeats. At the end of each simulation, each cell position with respect to the vessel longitudinal direction (y-axis) was calculated as the average among the y-coordinate of that specific cell memAgents.

We evaluated Alk1 KO tip cell competition by creating mosaic vessels with 5 wild type and 5 mutant cells in random initial positions and quantifying the average position of WT and mutant cells along the gradient. We evaluated the effect of Alk1 knockout on cell rearrangement by creating all-mutant vessels and counting the number of changes of cell order along the gradient. We first filtered the kymograph time series with a uniform filter of length 1 hour to remove short-lived position changes and then counted the number of cell rearrangements over 13000 timesteps (∼109 hours), comparing to an all-wild type vessel.

### ODE model of EC-pericyte crosstalk

#### Model assumptions and parameter choice

The ODE model of EC signaling was extended by adding the Notch signaling crosstalk of ECs with pericytes, as motivated by previous experiments (Benedito *et al*., 2009; Larrivée *et al*., 2012; Kakuda and Haltiwanger, 2017; Kakuda *et al*., 2020; Tefft *et al*., 2022). The cell-cell signaling schematic is presented in Fig. 5A. Here we highlight the assumptions that led to the model enhancement.

Pericytes were assumed to express Notch3, which could be activated by Jag1 in neighboring ECs (Benedito *et al*., 2009; Tefft *et al*., 2022). The interaction between Jag1 in ECs and Notch3 in pericytes was assumed to be equivalent to the interaction between Dll4 and Notch1, with thus the same model parameters. Notch3 activation following Jag1-Notch3 complex formation caused Dll4 expression in pericytes, in the same fashion as the Vegf-induced Dll4 expression in ECs. The Dll4 ligands that were expressed in pericytes could then bind and activate Notch1 present in neighboring ECs (Tefft *et al*., 2022), in the same way as the Dll4 expressed in ECs.

Bmp9 was assumed to initiate Jag1 expression in ECs (Larrivée *et al*., 2012), in the same order of magnitude as the VEGF-mediated expression of Dll4 in tip cells. In addition to the Notch3 in pericytes, Jag1 could also bind and activate Notch1 in ECs (Benedito *et al*., 2009). Notch1-Jag1 binding was assumed to occur at a comparable rate compared to Notch1-Dll4 binding, with this rate linearly proportional with LFng (Kakuda *et al*., 2020). Activation of Notch1 by Jag1 was assumed to occur at a much lower rate compared to Dll4-Notch1 activation, with the rate further decreased (10 times) by LFng, in a step-wise fashion (Kakuda *et al*., 2020). Finally, the Notch1-Jag1 complex dissociation rate was tuned to observe a considerable inhibiting effect of Jag1 for Notch signaling, in the case of EC monoculture with contemporaneous coating of both Jag1 and Dll4 (Benedito *et al*., 2009).

#### Simulations

Each EC was assumed to be in contact with one pericyte, and interacting via Notch signaling. *In vitro* qPCR experiments with EC monolayers and co-culture with pericytes were simulated by assuming that cells were exposed to insignificant Vegf (*V* = 0). The ODEs were solved with a final time equal to 48h, and external Bmp9 was added only 24h hours after the start of the experiment. The parameter values ℎ_*H*_*bmp*_ and ℎ_2_*bmp*_ quantifying Bmp9 effects on Hes1/Hey1 and LFng were left unchanged compared to the EC monoculture experiments. Jag1 expression without Bmp9 was assumed to be two orders of magnitude lower compared to the Bmp9-induced value. Jag1 siRNA was simulated by imposing an unchanged expression despite Bmp9 injection. LFng siRNA was simulated analogously.

EC stabilization occurring at the end of angiogenesis was simulated by adopting the same ODE model. For the first 24h, only ECs were simulated, exposed to VEGF (*V*_*ext*_ = 0.04 units) to obtain an initial tip/stalk pattern. For the following 24h, as in the simulation of tip/stalk pattern formation, Bmp9 injection was modeled by increasing the basal expression of Hes1/Hey1 and LFng by four times (ℎ_*H*_*bmp*_ = 4, and ℎ_2_*bmp*=_ = 4). In addition, Bmp9 induced Jag1 expression in the same fashion as in the qPCR experiments, and pericytes were added to simulate the Jag1-mediated EC-pericyte signaling crosstalk. The time necessary to stabilize ECs was evaluated in the same way as for the tip/stalk pattern formation. Endothelial cells were assigned a tip cell phenotype if *f*_*min*_ > 25 *c*. *u*. and a stalk cell phenotype otherwise. The vessel was assumed to be stabilized when all ECs exhibited a stalk cell phenotype. For an approximation of the rate of stabilization, the time necessary to obtain all stalk cells with different conditions was normalized over the time obtained with the original parameters, with Bmp9 and pericyte addition.

## Acknowledgements

Funding: this study was supported by the Marie Sklodowska-Curie Global Fellowship, grant number 846617 (to TR), and by the research program NWO Rubicon, which is (partly) financed by the Dutch Research Council (NWO), with project number 019.183EN.025 (to TR). MU was supported by an EMBO long-term fellowship (EMBO ALTF811-2018), the Center for Multiscale & Translational Mechanobiology at Boston University, and an AHA postdoctoral fellowship (828475). MU and CSC were supported by NIH (EB00262 and HL147585). SPH was supported by the Wellcome Trust (219500/Z/19/Z) and the British Heart Foundation (PG/18/67/33891). BL was supported by the Canadian Institutes of Health Research (FRN 363540) and the Natural Sciences and Engineering Research Council of Canada (RGPIN/05222-2018). KB, PO and IMA were supported by the Francis Crick Institute, which receives its core funding from Cancer Research UK (FC001751), the UK Medical Research Council (FC001751), and the Wellcome Trust (FC001751).

## Author contributions

Conceptualization: TR, BL, KB. Methodology: TR, RT, SPH, BL, KB. Software: TR, PQ. Validation: TR, EH, KL, KN, BL. Formal Analysis: TR, RT, EH, BL. Investigation: TR, RT, EH, KL, KN, IMA, MU. Writing – Original Draft: TR. Writing – Review & Editing: RT, EH, PQ, KL, KN, IMA, MU, SPH, CSC, BL, KB. Visualization: TR, BL, KB. Supervision: CSC, SPH, BL, KB. Project Administration: TR, BL, KB. Funding Acquisition: TR, SPH, CSC, BL, KB.

