## Supplementary Information for "Bmp9 regulates Notch signaling and the temporal dynamics of angiogenesis via Lunatic Fringe"

[1]Department of Biomedical Engineering, Eindhoven University of Technology, Eindhoven, 5612 AZ, the Netherlands
[2]Institute for Complex Molecular Systems (ICMS), Eindhoven University of Technology, Eindhoven, 5612 AZ, the Netherlands
[3]The Biological Design Center and Department of Biomedical Engineering, Boston University, Boston, MA 02215, USA
[4]Faculty of Biology, Medicine & Health, The University of Manchester, Manchester, M13 9PL, United Kingdom
[5]Maisonneuve-Rosemont Hospital, Montreal, QC H1T 2M4, Canada
[6]The Francis Crick Institute, London, NW1 1AT, UK
[7]Department of Ophthalmology, Université de Montréal, Montreal, QC H3C 3J7, Canada
[8]The Wyss Institute for Biologically Inspired Engineering, Harvard University, Boston, MA 02115, USA
[9]Department of Informatics, King's College London, London, WC2B 4BG, UK
[10]Lead author
[†]Contributed equally

### Supplementary Figure 1

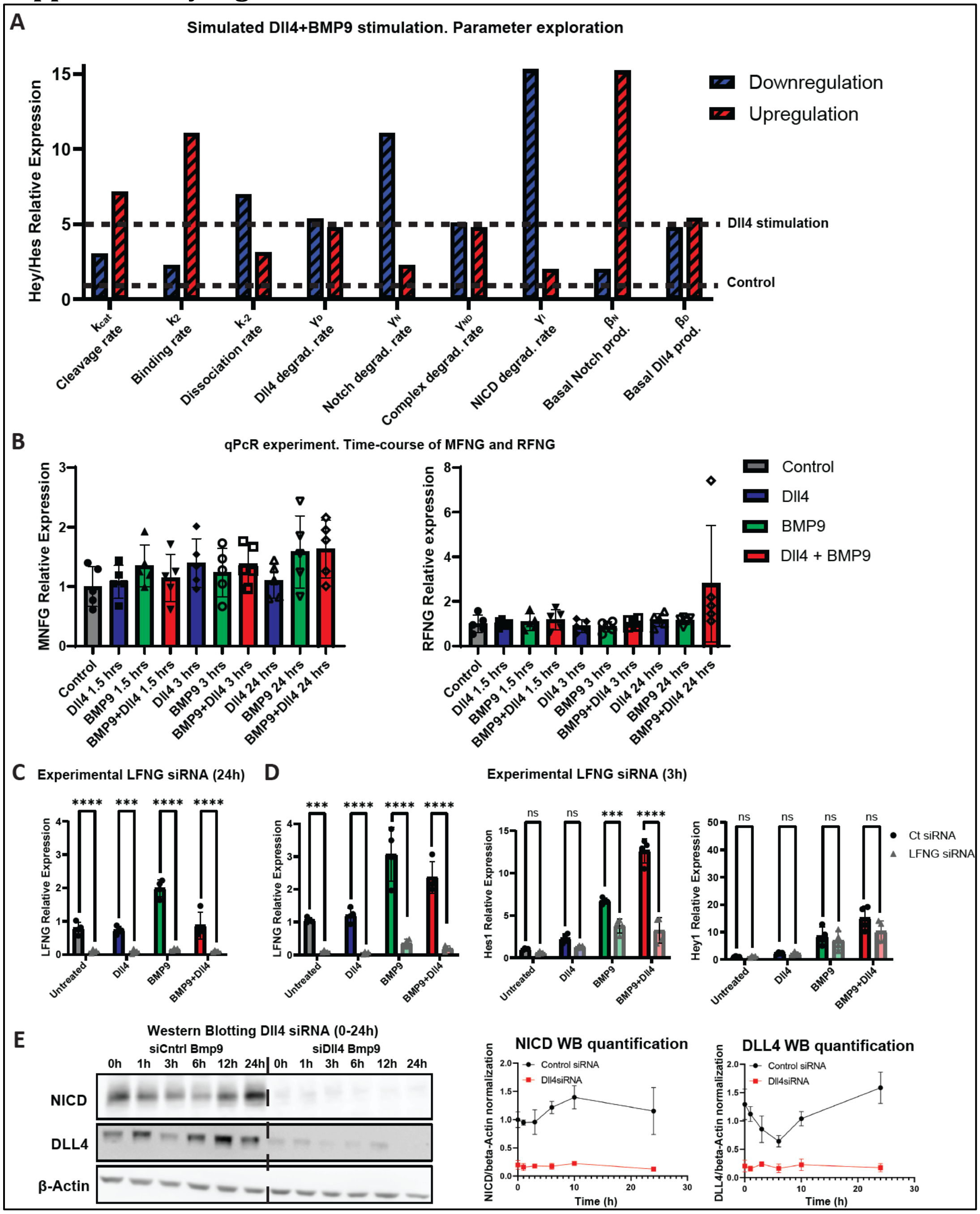


*Supplementary figure 1*

*(A) Computational analysis of the expression of Hey1/Hes1 in HUVECs exposed to Dll4 coating for 24h. The results of the simulations with original parameters, with and without Dll4 coating, are reported with horizontal dashed lines. Each bar corresponds to a different simulation obtained by doubling (red) or halving (blue) the model parameters reported in the horizontal axis.*

*(B-D) qPCR analysis of MFng, RFng, LFng, Hes1, Hey1 in HUVECs, induced by Bmp9 injection and/or Dll4 coating after 24h (B-C) or 3h (D) stimulation, with (bright color) or without (shaded color) LFng siRNA treatment. Number of repeats was higher than 4 in all cases ((B) N=5; (C-D) N=4).*

*(E) Western blot analysis of NICD and Dll4 compared to the safe-keeping gene beta-actin in HUVECs with control (left) or Dll4 siRNA treatment (right) upon stimulation with 10ng/ml Bmp9, up to 24h, with relative quantification normalized over unstimulated data (graphs). The number of independent repeats per condition was N = 3.*

*All values are mean +/- SD. *p < 0.05; **p<0.01; ***p<0.001; ****p<0.0001; ANOVA.*

### Supplementary Figure 2

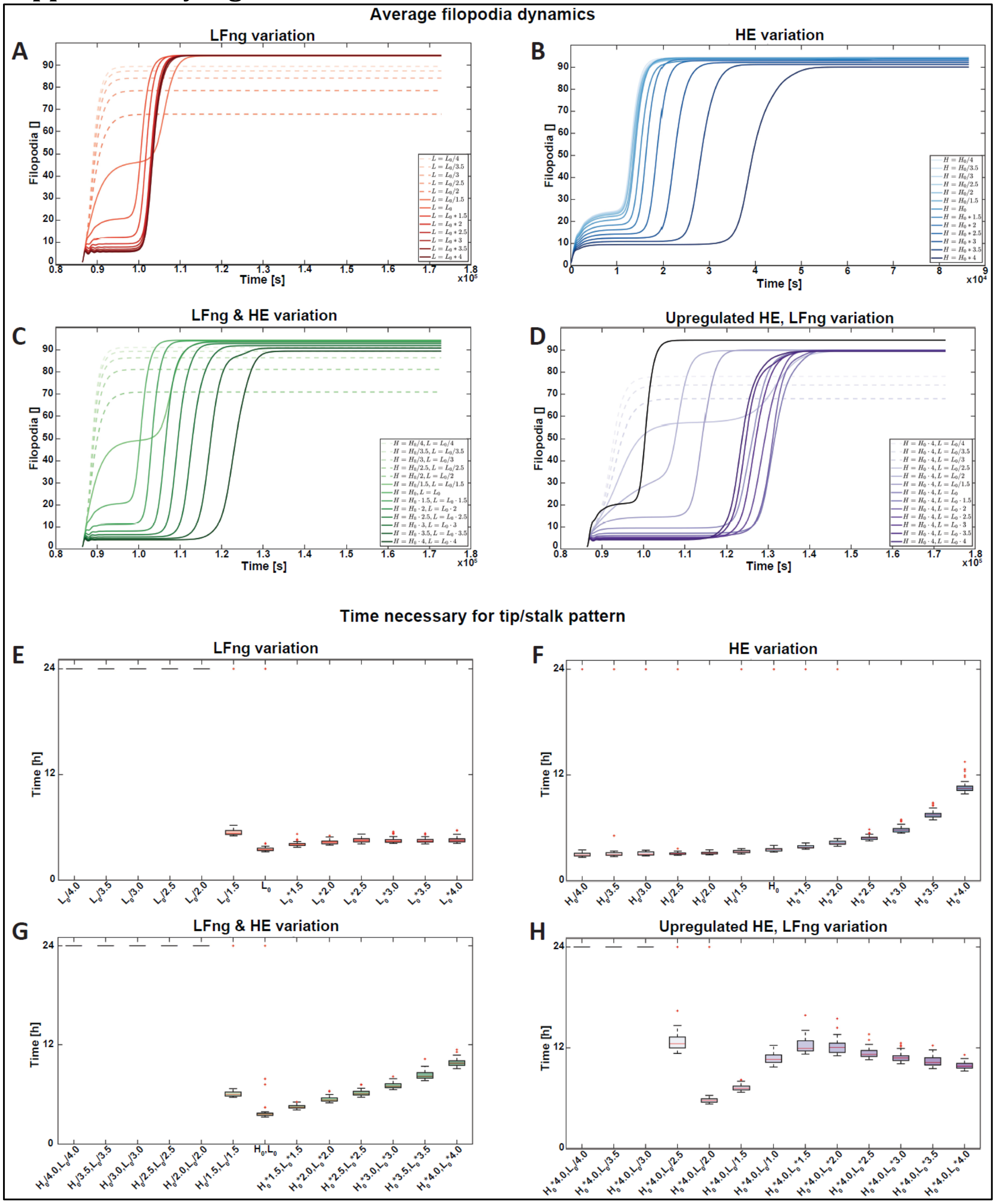


*Supplementary figure 2: temporal dynamics of tip/stalk selection affected by Hes1/Hey1 and LFng changes. (A-D) Computational analysis of the average amount of filopodia predicted for a row of 10 HUVECs exposed to no Vegf for 24h, followed by Vegf exposure for 24h. Each line represents a different parameter set obtained with the parameter modifications present in the captions. Dashed lines indicate that the tip/stalk pattern was not achieved for those sets of parameters.*
*(B) Computational analysis of the time necessary to obtain the tip/stalk pattern for a row of 10 HUVECs exposed to no Vegf for 24h, followed by Vegf exposure for 24h. Time = 0h was chosen to correspond with the initial time of Vegf exposure. Simulations that did not predict any tip/stalk pattern formation correspond to time to pattern values equivalent to 24h. In the boxplots, the central mark represents the median value. The boxes of the boxplots represent the quartiles of the distributions (25th-75th percentiles), with the whiskers extending to all the data not considered outliers, indicated with "+".*

### Supplementary Information. Methods

#### Computational model of EC signaling

A previous computational model (Venkatraman, Regan and Bentley, 2016) was extended to model the crosstalk of Vegf, Notch, and filopodia formation among a row of endothelial cells. The same assumptions as in the previous model (Venkatraman, Regan and Bentley, 2016) were taken, although the model was adapted to separately consider the proteins present in cell edges in contact with different cell neighbours. In particular, since a row of cells was considered, the ensemble of Notch receptors and ligands was divided into two groups; proteins on the left cell edge, indicated with the subscript $L$, and proteins on the right cell edge, with subscript $R$. Notch proteins were considered to diffuse between the two edges. In addition to this, the original model was modified to account for the Bmp9-mediated upregulation of both Hes1/Hey1 expression and the Notch receptor-ligand binding rate, modeled with a linear relationship. Finally, additional equations and terms were added to model the possible Notch activation as a result of ECs exposed to external Dll4 coating. Below the model equations are briefly summarized and motivated. In each equation, the subscript $i \in \mathbb{N}$ indicates a specific cell on the cell row, with $i-1$ and $1+1$ its neighbors on the left and right, respectively.

In the model, the Vegfr2 receptors presents in the EC $i$ are exposed to a level of Vegf $V_i$ dependent on the cell filopodia $f_i$:

$$V_i = (1 + k_3 f_i^{\,2}) V_0 \qquad (1)$$

where $V_0$ is an amount of reference of Vegf and the parameter $k_3$ scales the influence of filopodia. Vegf can bind to Vegfr2 ($R_i$) and form a Vegfr2-Vegf complex ($R.V_i$) which increase filopodia formation, while the expression of Vegfr2 decreases in response to Hes1/Hey1 ($H_i$) and degradation as follows:

$$\frac{df_i}{dt} = \beta + k_f V.R_i - k_{-f} f_i \qquad (2)$$

$$\frac{dR_i}{dt} = \gamma - k_1 V_i R_i + k_{-1} V.R_i - k_{inh} R_i H_i^2 - \phi R_i \qquad (3)$$

$$\frac{dV.R_i}{dt} = k_1 V_i R_i - k_{-1} V.R_i - \phi V.R_i \qquad (4)$$

where $\beta$ and $\gamma$ are the basal filopodia formation and production rate of filopodia and Vegfr2, respectively; $k_f$ scales the filopodia formation in response to Vegfr2 activation; $k_{-f}$ represents the turnover rate of filopodia; $k_1$ and $k_{-1}$ are respectively the protein association and dissociation rates; $\phi$ labels the protein degradation rate; and $k_{inh}$ scales the inhibition of Vegfr2 by Hes1/Hey1.

Vegfr2 activation via Vegf, quantified by $V.R_i$, leads to downregulation of Dll4 ($D_i$) expression, which can in turn bind and activate to Notch1 ($N_i$) present in neighboring cells. Notch1 can be activated also by external Dll4 ($D_{ext}$), forming associated complexes ($D_{ext}.N_i$). The content of Dll4, Notch1, and their complex in the left edge of each cell $i$ then varies according to:

$$\frac{dD_{L_i}}{dt} = \frac{1}{2}\left(\beta + \theta \frac{V.R_i^2}{1+V.R_i^2}\right) - 2k_{2\_bmp} D_{L_i} N_{R_{i-1}} + k_{-2} D.N_{R_{i-1}} - \phi D_{L_i} + W\left(\frac{D_L + D_R}{2} - D_L\right) \qquad (5)$$

$$\frac{dN_{L_i}}{dt} = \frac{\gamma}{2} - 2k_{2\_bmp} N_{L_i}\left(D_{R_{i-1}} + D_{ext}\right) + k_{-2}\left(D.N_{L_i} + D_{ext}.N_{L_i}\right) - \phi N_{L_i} + W\left(\frac{N_L + N_R}{2} - N_L\right) \qquad (6)$$

$$\frac{dD.N_{L_i}}{dt} = 2k_{2\_bmp}N_{L_i}D_{R_{i-1}} - k_{-2}D.N_{L_i} - \phi D.N_{L_i} - k_{cat}D.N_{L_i} \quad (7)$$
$$\frac{dD_{ext}.N_{L_i}}{dt} = 2k_{2\_bmp}N_{L_i}D_{ext} - k_{-2}D_{ext}.N_{L_i} - \phi D_{ext}.N_{L_i} - k_{cat}D_{ext}.N_{L_i} \quad (8)$$

The time variations of the proteins on the right, indicated with subscript $R$, are analogous. Here, $\theta$ scales the downregulation of Dll4 in response to Vegfr2 activation; $k_{2\_bmp}$ and $k_{-2}$ are respectively the dissociation and association rates of Dll4 and Notch1; $W$ scales the movement rate of unbound Notch proteins across the two cell edges; and $k_{cat}$ labels the Notch1 activation rate once complexes with either cellular or extracellular Dll4 are formed. Note that, compared to the original model, the Notch protein production and binding rate were scaled by a factor 2, to accommodate for the two cell edges. The Bmp9-mediated upregulation of the Notch receptor-ligand binding rate was modeled linearly as follows:

$$k_{2\_bmp} = h_{2\_bmp}k_2 \quad (9)$$

where $k_2$ is the homeostatic binding rate and $h_{2\_bmp}$ linearly scales Bmp9 effects.

Notch activation leads to an increase in Notch intracellular domain ($I_i$) and a consequential increase in Hes1/Hey1 ($H_i$) expression as follows:

$$\frac{dI_i}{dt} = k_{cat}\left(D.N_{L_i} + D.N_{R_i} + D_{ext}.N_{L_i} + D_{ext}.N_{R_i}\right) - \phi I_i \quad (10)$$
$$\frac{dH_i}{dt} = \beta_{H\_bmp} + \theta\frac{I_i^2}{1+I_i^2} - \phi I_i. \quad (11)$$

where $\beta_{H\_bmp}$ is the Hes1/Hey1 expression as affected by Bmp9:

$$\beta_{H\_bmp} = h_{H\_bmp}\beta \quad (12)$$

where $\beta$ indicates the basal production rate, linearly scaled by $h_{H\_bmp}$ as a result of Bmp9.

The parameters were left unchanged compared to the original model implementation (Venkatraman, Regan and Bentley, 2016). The new parameters describing the effects of external Dll4 ($D_{ext}$) and Bmp9 ($k_{2\_bmp}$ and $\beta_{H\_bmp}$) were calibrated against previous experiments (Larrivée *et al.*, 2012). Finally, for simplicity, the unbound Notch protein movement rate ($W$) was kept high enough to ensure fast protein homogenization. The model parameters are listed in Table S1, where "cu" stands for concentration units.

**Computational model of EC-pericyte signaling**

Briefly, Notch signaling between ECs and pericytes was modeled by adding Jag1 in ECs, which could bind and activate Notch1 in ECs and Notch3 in pericytes. Notch3 activation led to Dll4 expression in pericytes, which in turn could bind and activate Notch1 in ECs. More details on the assumptions of the model as motivated by previous experiments are reported in the main text. Here, we report the equations of the extended Notch model. Again, the ensemble of Notch receptors and ligands was divided into two groups, one for each cell edge. We describe here the equations for the left edge with subscript $L$. Analogous equations were used for the right edge. The equations for Vegf signaling and filopodia formation, as well as the equations of Dll4 and Hes1/Hey1 in ECs, and Bmp9 effects on LFng are not reported, since Eqs. 1-5, 8, and 10-12 Were left unchanged. The remaining equations are described highlighting the changes compared to the EC signaling model alone.

Bmp9 induced Jag1 ($J_i$) expression in ECs, at a rate linearly dependent on the amount of Bmp9:

$$\beta_{J\_bmp} = h_{J\_bmp} 1 \tag{13}$$

Jag1 ($J_i$) could bind to: Notch1 ($N_i$) in neighboring ECs, forming protein complexes $J_i.N_i$ at a rate $k_{2\_bmp}$ influenced by Bmp9 via LFng; Notch3 ($N_{3i}$) in neighboring pericytes, forming protein complexes $J_i.N_{3i}$ at a rate independent from Bmp9 ($k_2$). These protein complexes can dissociate at a rate $k_{-2J}$ and $k_{-2}$, respectively, such that the time variation of Jag1 present on the left edge of the cell $i$ can be described as:

$$\frac{dJ_{L_i}}{dt} = \frac{\beta_{J_{bmp}}}{2} - k_{2_{bmp}} J_{L_i} N_{R_{i-1}} + k_{-2J} J.N_{R_{i-1}} - k_2 J_{L_i} N_{3i} + k_{-2} J.N_{3L_i} - \phi J_{L_i} + W\left(\frac{J_L + J_R}{2} - J_L\right) \tag{14}$$

while the Notch3 content in pericytes varies over time according to:

$$\frac{dN_{3_i}}{dt} = \frac{\gamma}{2} - 2k_2 N_{3_i}\left(J_{R_i} + J_{L_i}\right) + k_{-2}\left(J.N_{3R_i} + J.N_{3L_i}\right) - \phi N_{3_i} \tag{15}$$

Upon protein complex formation, Notch3 is activated and leads to an increase in Notch3 intracellular domain ($I_{3_i}$). This upregulates pericyte expression of Dll4 ($P_i$), which can bind in turn to Notch1 in neighboring ECs forming Dll4-Notch1 protein complexes at the interface between ECs and pericytes ($P.N_{L_i}$):

$$\frac{dI_{3i}}{dt} = k_{cat}\left(J.N_{3L_i} + J.N_{3R_i}\right) - \phi I_{3i} \tag{16}$$

$$\frac{dP_i}{dt} = \frac{1}{2}\left(\beta + \theta \frac{I_{3i}^2}{1 + I_{3i}^2}\right) - 2k_{2_{bmp}} P_i N_{R_{i-1}} + k_{-2}(P.N_{R_{i-1}} + P.N_{L_{i-1}}) - \phi P_i \tag{17}$$

Notch1 in ECs can be activated by Dll4 ($D_i$) and Jag1 in ECs, as well as Dll4 ($P_i$) from pericytes, with which it forms protein complexes $P_i.N_i$. The interaction between Notch1 and Dll4 is always the same, irrespective of the cell source, such that the variation of Notch1 and related complexes can be tracked via the following ODEs:

$$\frac{dN_{L_i}}{dt} = \frac{\gamma}{2} - 2k_{2\_bmp} N_{L_i}\left(D_{R_{i-1}} + P_i\right) + k_{-2}\left(D.N_{L_i} + P.N_{L_i}\right) - k_{2\_bmp} N_{L_i} J_{R_{i-1}} + k_{-2J} J.N_{L_i} - \phi N_{L_i} + W\left(\frac{N_L + N_R}{2} - N_L\right) \tag{18}$$

$$\frac{dD.N_{L_i}}{dt} = 2k_{2\_bmp} N_{L_i} D_{R_{i-1}} - k_{-2} D.N_{L_i} - \phi D.N_{L_i} - k_{cat} D.N_{L_i} \tag{19}$$

$$\frac{dJ.N_{L_i}}{dt} = k_{2\_bmp} N_{L_i} J_{R_{i-1}} - k_{-2J} J.N_{L_i} - \phi J.N_{L_{i_i}} - k_{cat\_LF} J.N_{L_i} \tag{20}$$

$$\frac{dP.N_{L_i}}{dt} = 2k_{2\_bmp} N_{L_i} P_i - k_{-2} P.N_{L_i} - \phi P.N_{L_i} - k_{cat} P.N_{L_i} \tag{21}$$

The parameter $k_{cat\_LF}$ scales the activation of Notch1 by Jag1 upon binding, and it's assumed to be dependent on the amount of LFng in a step-wise fashion. Given that LFng was not modeled explicitly but only via its effects on Notch1 receptor-ligand affinity as quantified by the parameter $h_{2\_bmp}$, the same parameter was used for the effects of LFng on Jag1-mediated Notch1 activation:

$$k_{cat\_LF} = \begin{cases} k_{cat}/10, & h_{2_{bmp}} < h_{min} \\ k_{cat}/100, & h_{2_{bmp}} \geq h_{min} \end{cases} \tag{22}$$

Finally, Notch1 activation led to an increase in Notch1 intracellular domain ($I_i$) as follows:

$$\frac{dI_i}{dt} = k_{cat}\left(D.N_{L_i} + D.N_{R_i} + P_i.N_{L_i} + P_i.N_{R_i}\right) + k_{cat\_LF}\left(J.N_{L_i} + J.N_{R_i}\right) - \phi I_i \qquad (23)$$

The parameters already present in the EC signaling model were left unchanged. The values of the remaining new parameters are reported in Table S2.

**Table S1. Parameter values of EC signaling model**

| Param. | Value | Description |
|---|---|---|
| $\boldsymbol{V_0}$ | 0.05 cu | Vegf concentration |
| $\boldsymbol{\gamma}$ | $5\cdot10^{-3}$ cu.sec$^{-1}$ | Production rate of Notch and Vegfr2 |
| $\boldsymbol{\beta}$ | $10^{-3}$ cu.sec$^{-1}$ | Basal formation/expression of filopodia/Dll4 and Hes1Hey1 |
| $\boldsymbol{h_{H\_bmp}}$ | 0 - 4 [] | Scaling factor of Bmp9 effects on Hes1/Hey1 |
| $\boldsymbol{\theta}$ | $10^{-1}$ sec$^{-1}$ | Scaling factor of Vegfr2/Notch effects on Dll4/Hes1Hey1 |
| $\boldsymbol{\phi}$ | $5\cdot10^{-3}$ sec$^{-1}$ | Degradation rate of |
| $\boldsymbol{k_1}$ | $10^{-1}$ cu$^{-1}$sec$^{-1}$ | Binding rate of the Vegf-Vegfr2 complex |
| $\boldsymbol{k_{-1}}$ | $10^{-3}$ sec$^{-1}$ | Dissociation rate of the Vegf-Vegfr2 complex |
| $\boldsymbol{k_2}$ | $10^{-3}$ cu$^{-1}$sec$^{-1}$ | Binding rate of the Notch1-Dll4 complex |
| $\boldsymbol{h_{2\_bmp}}$ | 0 - 4 [] | Scaling factor of Bmp9 effects on $k_2$ |
| $\boldsymbol{k_{-2}}$ | $10^{-1}$ sec$^{-1}$ | Dissociation rate of the Notch1-Dll4 complex |
| $\boldsymbol{k_3}$ | $5\cdot10^{-3}$ cu$^{-2}$ | Scaling factor of filopodia-Vegf positive correlation |
| $\boldsymbol{k_f}$ | $10^{-1}$ sec$^{-1}$ | Filopodia formation rate in response to Vegfr2 |
| $\boldsymbol{k_{-f}}$ | $10^{-3}$ sec$^{-1}$ | Filopodia turnover rate |
| $\boldsymbol{k_{inh}}$ | $5\cdot10^{-3}$ cu$^{-2}$sec$^{-1}$ | Scaling factor of Hes1/Hey1-Vegfr2 negative correlation |
| $\boldsymbol{k_{cat}}$ | $10^{-1}$ sec$^{-1}$ | Cleavage rate of the Notch1-Dll4 complex |
| $\boldsymbol{W}$ | $10^{-3}$ sec$^{-1}$ | Movement rate of unbound Notch proteins |

**Table S2. Additional values of EC-pericyte signaling model**

| Param. | Value | Description |
|---|---|---|
| $\boldsymbol{h_{J\_bmp}}$ | 0.01 cu.sec$^{-1}$ | Reference production rate of Jag1 in response to Bmp9 |
| $\boldsymbol{k_{-2j}}$ | $5\cdot10^{-4}$ sec$^{-1}$ | Dissociation rate of the Notch1-Jag1 complex |
| $\boldsymbol{h_{min}}$ | 0.25 [] | Threshold value of LFng effects on Notch1-Jag1 activation |

**Table S3. List of antibodies**

| Antibodies | Company | Cat. no. | Clonality | Clone no. | Dilution |
|---|---|---|---|---|---|
| LFNG | Cell Signaling | 66472 | Rabbit polyclonal | D6V2V | 1:100 (IHC) 1:1000 (WB) |
| Cleaved Notch1 | Cell Signaling | 4147 | Rabbit polyclonal | D3B8 | 1:100 |
| β-actin | Santa Cruz | sc-47778 | Mouse monoclonal | C4 | 1:2000 |
| Dll4 | Cell Signaling | 96406 | Rabbit polyclonal | D7N3H | 1:1000 |

**Supplementary Video 1:** Representative time-lapse confocal movie of EC nuclei dynamics in intersegmental vessel (ISV) sprouts in control (MOC) Tg(kdrl:nlsEGFP)$^{zf109}$ embryos (from Fig. 3F).

**Supplementary Video 2:** Representative time-lapse confocal movie of EC nuclei dynamics in intersegmental vessel (ISV) sprouts in lfng-knockdown Tg(kdrl:nlsEGFP)$^{zf109}$ embryos (from Fig. 3F).
